# Host genetic variants regulates CCR5 expression on immune cells: a study in people living with HIV and healthy controls

**DOI:** 10.1101/2022.06.19.496757

**Authors:** Jéssica C. dos Santos, Zhenhua Zhang, Louise E. van Eekeren, Ezio T. Fok, Nadira Vadaq, Lisa van de Wijer, Wouter A. van der Heijden, Valerie A. C. M. Koeken, Hans J.P.M. Koenen, Musa Mhlanga, Mihai G. Netea, André J. van der Ven, Yang Li

**Affiliations:** Department of Internal Medicine and Radboud Center for Infectious Diseases, Radboud University Medical Center, 6525 HP Nijmegen, the Netherlands; Department of Genetics, University of Groningen, University Medical Center Groningen, 9700 RB Groningen, the Netherlands; Epigenomics & Single Cell Biophysics Group, Department of Cell Biology, Radboud University, 6525 GA Nijmegen, the Netherlands; Laboratory Medical Immunology, Department of Laboratory Medicine, Radboud University Medical Center, 6525 HP Nijmegen, the Netherlands; Department of Human Genetics, Radboud University Medical Center, 6525 GA Nijmegen, the Netherlands; Radboud Institute for Molecular Life Sciences, 6525 GA Nijmegen, the Netherlands; Department of Immunology and Metabolism, Life and Medical Sciences Institute, University of Bonn, Germany; TWINCORE, Centre for Experimental and Clinical Infection Research, a joint venture between the Hannover Medical School and the Helmholtz Centre for Infection Research, Hannover, Germany; Department of Computational Biology for Individualised Medicine, Centre for Individualised Infection Medicine (CiiM), a joint venture between the Helmholtz-Centre for Infection Research (HZI) and the Hannover Medical School (MHH), Hannover, Germany

## Abstract

C-C chemokine receptor 5 (CCR5) is the main HIV co-receptor affecting susceptibility and disease course. Quantitative trait loci (QTL) mapping analysis was performed to assess genetic variants associated with CCR5 expression on circulating immune cells in 209 PLHIV using ART and 304 healthy controls, all of Western European ancestry. The proportions of CCR5 positive cells and CCR5 mean fluorescence intensity (MFI) were assessed by flow cytometry in monocytes and CD4^+^ and CD8^+^ T cell subsets using flow cytometry. We identified the rs60939770, which is an intergenic variant in *cis*-region to *CCR5* gene not in linkage disequilibrium with *CCR5d32,* related to the proportion of CCR5^+^ memory T regulatory cells, both in PLHIV and healthy controls. Two genome-wide significant loci, in linkage equilibrium with *CCR5d32*, were found to be associated with CCR5 MFI of multiple subsets of mostly differentiated memory T cells in both groups. The expression of nearby chemokines receptors (*CCR1*, *CCR2*, *CCR3, CCRL2*), existing in the same the same topologically associating domain, were also influenced by these genetic variants. Furthermore, we show the genetic variants which modulate CCR5 surface expression affect the production of other inflammatory mediators, including monocyte- and lymphocyte-derived cytokines as well as CCL4 and IL-8. Our data indicate that the genetic regulation of CCR5 expression is cell-specific and affects the production of various inflammatory mediators.

**Author Summary:** CCR5 plays a important role in the acquisition of HIV and it is associated to immune activation in people living with HIV (PLHIV). Using samples of cohorts composed of healthy individuals and PLHIV, we sought to map genomic regions that influence CCR5 expression on monocytes and subsets of CD4^+^ and CD8^+^ cells. We identified distinct genetic variants that are associated with CCR5 cell proportions or mean fluorescence intensity in subpopulations of T cells with memory functions in both healthy and PLHIV. The genetic variants also influenced the expression of other nearby chemokine receptors and the production of inflammatory mediators. Thus, we demonstrated that the genetic regulation of CCR5 expression is cell-type specific and may impact HIV susceptibility and disease progression.

## 1. Introduction

CD4 receptor-mediated entry of human immunodeficiency virus-1 (HIV-1) requires binding to C-C chemokine receptor 5 (CCR5) or C-X-C chemokine receptor 4 (CXCR4) as a co-receptor (1). While CCR5 is the main co-receptor for HIV-1 entry, its expression levels on the surface of specific CD4+ T cell populations have been shown to be associated with the response to treatment and disease progression in HIV infection (2). CCR5 ligands (CCL3, CCL4, CCL5 and CCL3L1) also play an important role in innate and adaptive immune responses, further highlighting the importance of CCR5 and its downstream signaling. As such, dysregulated CCR5 expression could contribute to non-AIDS associated comorbidities that become more prevalent in people living with HIV (PLHIV), despite highly effective long-term suppressive combination antiretroviral treatment (ART) (3).

Many studies have been identified genetic factors that influence HIV acquisition and progression (1). A consistent finding is a 32-base pair deletion in the open reading frame (ORF) of the CCR5 gene, resulting in a defective CCR5 that cannot emerge on the surface of the cell after translation (referred to as *CCR5d32* (rs333)). Heterozygous *CCR5d32* individuals have reduced surface levels of CCR5 allowing PLHIV to benefit from slower disease progression, while the complete absence of CCR5 on the cell surface due to a homozygous *CCR5d32* deletion can prevent infection by CCR5-tropic strains of HIV (4). Moreover, stem cell transplantation from a homozygous *CCR5d32* donor has led to functional cure of HIV in the infected recipient (5). These consequences of *CCR5d32* highlight the possible clinical impact of other genetic variants on CCR5 expression levels.

Previous studies have directly evaluated CCR5 genotypes in relation to HIV pathogenesis and overlooked genome-wide genetic variations associated with CCR5 expression on immune cells targeted by HIV-1 (6, 7). These genotypes include specific single nucleotide polymorphisms (SNPs) in the CCR5 and CCR2 coding region that were grouped into seven phylogenetically distinct clusters known as the CCR5 human haplotypes (HH) A-G (6). The current evidence on the association between specific CCR5 haplotypes and CCR5 expression is scarce and contradictory (8, 9). Moreover, the link between genetics and CCR5 expression have not been studied for specific HIV-1 relevant cell subsets at different stages of differentiation. The latter is particularly important as terminally differentiated memory immune cells express higher levels of CCR5 than their naïve counterparts (10). This increase of CCR5 expression on CD4^+^ T cells with memory functions has been described as a marker of disease progression, since optimal conditions for viral replication are provided by these subsets of cells (11). Thus, although several SNPs related to HIV pathogenesis are found in the CCR5 gene, it remains unknown whether genome-wide genetic variations on specific cell-types determine interindividual CCR5 expression levels in PLHIV with an intact CCR5 ORF.

In this study we aimed to conduct a in a genome-wide association study (GWAS) PLHIV to assess the contribution of host genetic variation on the cell surface expression of CCR5 and the proportion of CCR5-expressing circulating immune cell subsets. As one of the GWAS SNPs (rs11574435) mapped to the intronic region of a previously described, antisense transcribed sequence named *CCR5AS* (12), we also aimed to assess its effects more broadly by measuring*, CCR5* as well as *CCR5AS* and nearby genes in PLHIV. Furthermore, our findings in PLHIV were corroborated in an independent cohort (300BCG cohort) of healthy individuals (13).

### 2. Results

### 2.1 Characteristics of the study populations

Two independent cohorts of adults (18 years and above) with Western European ancestry were included in this study. This consisted of 209 people living with HIV (PLHIV) using long-term ART (200HIV) and 304 healthy individuals (300BCG study) (Fig. 1A). The average age of PLHIV and the healthy individuals was 52 years and 23 years, respectively, of which 91% and 43% respectively were males. In the PLHIV, HIV transmission routes included homosexual contact (157/209), heterosexual contact (39/209), intravenous drug use (IDU, 3/209), needle stick injury (1/209), and contaminated blood products (1/209). For the remaining 8/209 participants, the route of transmission was unknown. The PLHIV have the following HIV-specific characteristics: CD4 nadir: median 250 10^6^ cells/L (IQR 230), latest CD4: median 660 10^6^ cells/L (IQR 330), zenith HIV-RNA: median 100.000 copies/ml (IQR 345.591), and cART duration: median 6.61 years (IQR 7.70). A total of 67% (139/209) used an integrase inhibitor, 30% (63/209) non-nucleoside analogue and 15% (32/209) a protease inhibitor. The HIV-RNA viral load was beneath the detection limit in 203/209 PLHIV.

**Fig 1.**
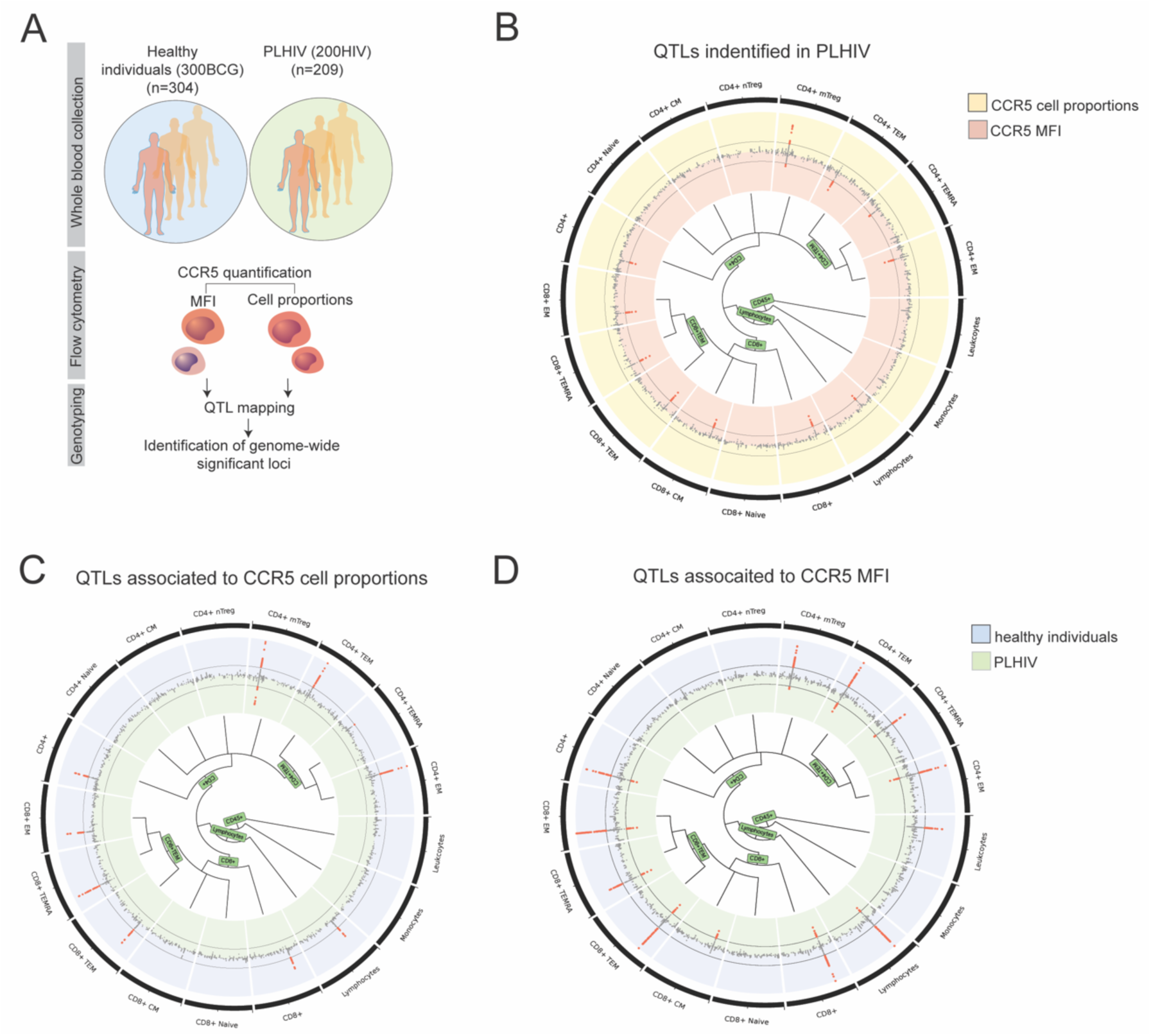
Genome-wide significant QTLs and their shared associations on multiple immune cell subpopulations. (A) The study design. (B), (C) and (D) are all combined Manhattan plots of multiple immune cell subpopulations (chromosome 3). (B) Includes associations for both CCR5 MFI (red) and cell proportions (CP) (yellow) for PLHIV. (C) Consists of associations for CP from PLHIV (green) and healthy individuals (blue) while (D) shows associations for MFI from PLHIV (green) and healthy individuals (blue). From outer to inner, the first track (black) shows assessed immune cell subpopulation name, where each sector represents one cell types; the second and third tracks include P-value Manhattan plots for each cell type assessed for CP (yellow) and MFI (red) in (B), or PLHIV (green) and healthy individuals (blue) in (C) and (D), respectively; the innermost is hierarchical tree of cell types.

### 2.2 Identifying genome-wide genetic determinants of CCR5 surface expression

To explore how genetics modulate the expression of CCR5 on the surface of different immune cell subsets, we conducted quantitative trait loci (QTL) mapping analysis using genotype data in the two independent cohorts. CCR5 expression on the surface of several subsets of immune cells was measured in both cohorts and expressed as the geometric mean of fluorescence intensity (MFI) and proportions of CCR5 positive cells (cell proportion (%)) (Fig. 1A, S1 Fig). The distributions of CCR5 MFI and proportions across the subpopulations of immune cells are shown in S3 Fig.

**Fig 3.**
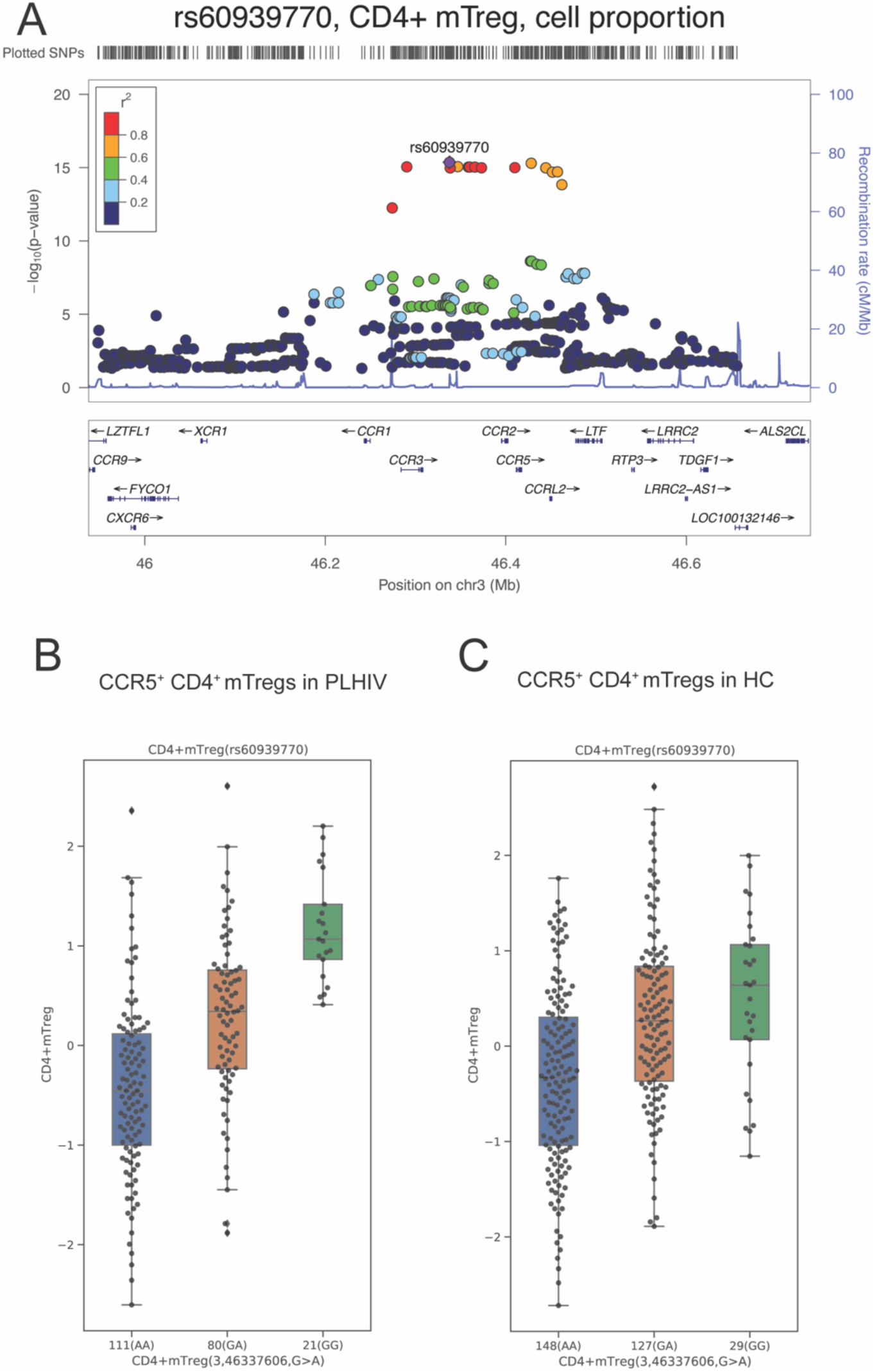
Genetic effect of the rs60939770 SNP on CCR5^+^CD4^+^ mTreg cells of PLHIV and healthy individuals. (A) Regional plot (LocusZoom) showing the QTL associated with CCR5 cell proportions in CD4^+^ mTreg of PLHIV, where the top SNP is rs60939770. (B) and (C) are boxplots of proportions of CCR5 positive CD4+ mTregs stratified according to the rs60939770 genotypes in PLHIV (P-value = 4.29 × 10^-16^) and HC (P-value = 3.18 × 10^-10^), respectively. Horizontal line in the boxplot, median; ends of the boxes, upper and lower quartiles.

After testing the association between common variants (MAF > 0.1) and CCR5 MFI or cell proportions using a linear regression model with age and sex corrected, we identified five independent genome-wide significant loci (p < 5 × 10^-8^) associated with CCR5 MFI or cell proportions in PLHIV (S1 Table). CCR5 MFI and cell proportions showed associations with three independent variants in the *cis-*region to the CCR5 locus (Fig. 1B) on chromosome 3. Moreover, two *trans*-loci variants associated with CCR5 MFI or cell proportions were located at chromosome 2 (S3 Fig). Interestingly, the majority of significant associations (75%) were found in relation to T cells with memory functions, suggesting the importance of genetics for CCR5 cell-surface expression in long-lasting populations of immune cells.

To validate these genetic associations and verify their specificity to PLHIV, we performed a QTL mapping analysis of the same measurements of the relevant immune cell subsets in an independent cohort of healthy controls (HC). We identified a common genetic loci that was associated with the proportions of CCR5^+^CD4^+^ mTregs for both PLHIV and HC (Fig. 1C). The genetic variant associated with CCR5 MFI in PLHIV was also identified in HC in the same subsets of CD4^+^ and CD8^+^ T cells, except for CD8^+^ central memory T cells in which the association was found in PLHIV only (Fig. 1D). Two additional genetic variants in the the *cis-*region of the CCR5 locus and one *trans*-loci variant were identified associated with both CCR5 MFI and cell proportions in subpopulations of CD4^+^ and CD8+ cells of HC only (S2 Table). Besides the validation of the common genetic associations in two independent cohorts, our results using both cohorts of PLHIV and HC allowed the identification of genetic variants that are important for CCR5 surface expression in specific immune cells subpopulations of PLHIV (Fig. 2). As CCR5 is the major co-receptor of HIV in immune cells (1), these genetic variants might have a relevant influence on the disease outcome in PLHIV and susceptibility of acquiring HIV in healthy controls.

**Fig 2.**
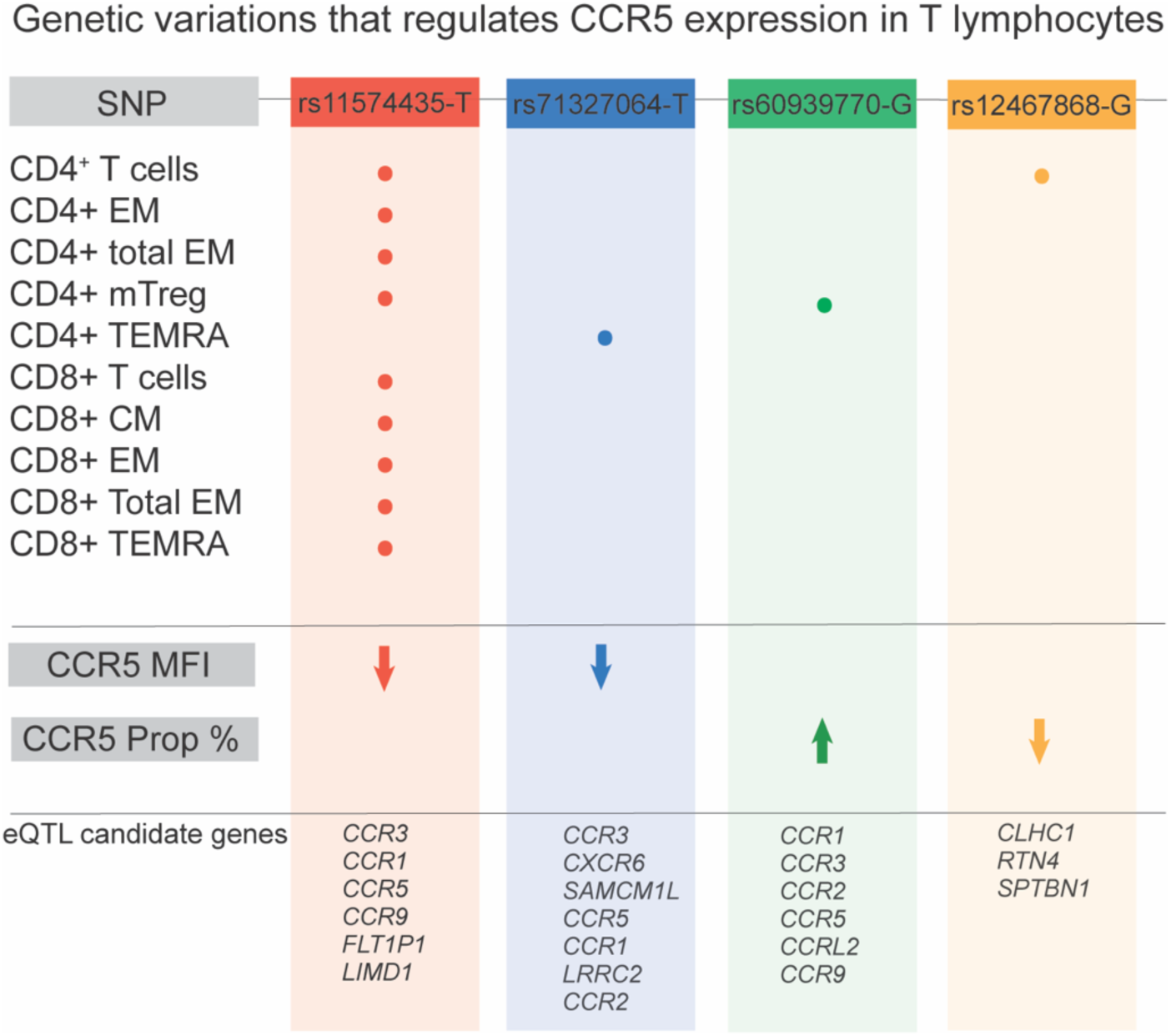
The loci representing the risk variants identified associated with geometric mean fluorescence intensity (MFI) and proportions of CCR5 positive cells or cell (prop %) in both PLHIV and HC. The arrow indicates the directionality of the changes in CCR5 MFI and prop % surface expression in individuals carrying the effect allele of each SNP in the indicated cell-type. Expression QTL results of candidate genes in whole blood of healthy individuals.

### 2.3 *Cis*- and *trans*-genetic effects on CCR5 proportions in CD4^+^ mTreg and total CD4^+^ T cells

The strongest association (*cis*-SNP rs60939770, chromosome 3, P-value = 4.29 × 10^-16^) was identified for the proportion of CCR5^+^CD4^+^ mTreg cells in PLHIV (Fig. 3A). Of importance, these effects were replicated in the HC cohort at genome-wide significance (P-value = 3.18 × 10^-10^). PLHIV carrying at least one rs60939770-G allele had a significantly higher proportion of CCR5^+^CD4^+^ mTreg cells than subjects homozygous for the A allele (Fig. 3B). The same genetic effect was also observed in the HC (Fig. 3C). The rs60939770 SNP is an intergenic variant in *cis*-region to the *CCR5* gene and it is in a linkage disequilibrium (LD) with the rs1015164 SNP (R^2^ = 0.6823, D’ = 0.8594, P-value < 0.0001). (Table S1). Our results reveal that the rs1015164 is strongly correlated to the the percentages of CD4^+^ mTreg cells expressing CCR5 in PLHIV and HC (P-value < 5 × 10^-8^) (S4A-C Fig). Individuals with the rs1015164-A was shown to have higher proportion of CCR5^+^CD4^+^ mTreg cells compared to rs1015164-G in both PLHIV and HC (S4B-D Fig). rs1015164 lies in the antisense long noncoding RNA *CCR5AS* and has previously been associated with increased CCR5 MFI surface expression in bulk memory CD4^+^ T cells and effector memory CD4^+^ T cells of healthy individuals (12). Given the previously reported effects of rs1015164 in CCR5 MFI expression, we tested the influence of rs1015164 on CCR5 MFI levels in our dataset. Due to the cellular subset resolution afforded by our study, we found CCR5 MFI levels to be most altered by this polymorphism in CD4^+^ mTreg cells (S4E-G Fig). The rs1015164-A was associated with higher CCR5 MFI expression than rs1015164-G in CD4^+^ mTreg cells of PLHIV and HC cells (P-value = < 0.05) (S4F-H Fig). Even though the rs60939770 and rs1015164 are in LD, we demonstrated that they have different effects on CCR5 epxression, which rs60939770 modulates CCR5 cell proportions and the rs1015164 affects both proportions of positive cells and MFI. Collectively, these results suggested that the expression of CCR5 in CD4^+^ mTreg cells of PLHIV and healthy controls is under *cis*-genetic regulation.

A *trans*-loci genetic variant, rs12467868 was also identified and associated with the percentage of CCR5^+^CD4^+^ T cells (P-value = 4.07 × 10^-8^) in PLHIV only (Fig. 2, S1 Table). The rs12467868 SNP lies in the intron 3 of the *RPS27* gene, a coding ribosomal protein gene implicated in viral replication of DNA and RNA viruses (14).

### 2.4 The genetic variants associated to CCR5 MFI are present in differentiated CD4^+^ and CD8^+^ T cell subsets

For CCR5 surface expression (measured as MFI), we identified a genetic association with rs11574435 SNP (P-value < 5 × 10^-8^) in the majority of both CD4^+^ and CD8^+^ T cell subsets of both PLHIV and HC (Fig. 2, S1 Table). In PLHIV, the rs11574435-CC genotype was associated with higher CCR5 MFI expression than individuals with TC genotypes in the subpopulations of CD4^+^ and CD8^+^ T cells (Fig. 4). Within the same locus, the rs71327064 SNP (P-value < 6.22 × 10^-9^) was identified to be associated with CCR5 MFI expression in CD4^+^ TEMRA cells (Fig. 2, S1 Table). Furthermore, these findings of both SNPs were replicated in the HC cohort with the same allelic direction (P-value < 0.05).

**Fig 4.**
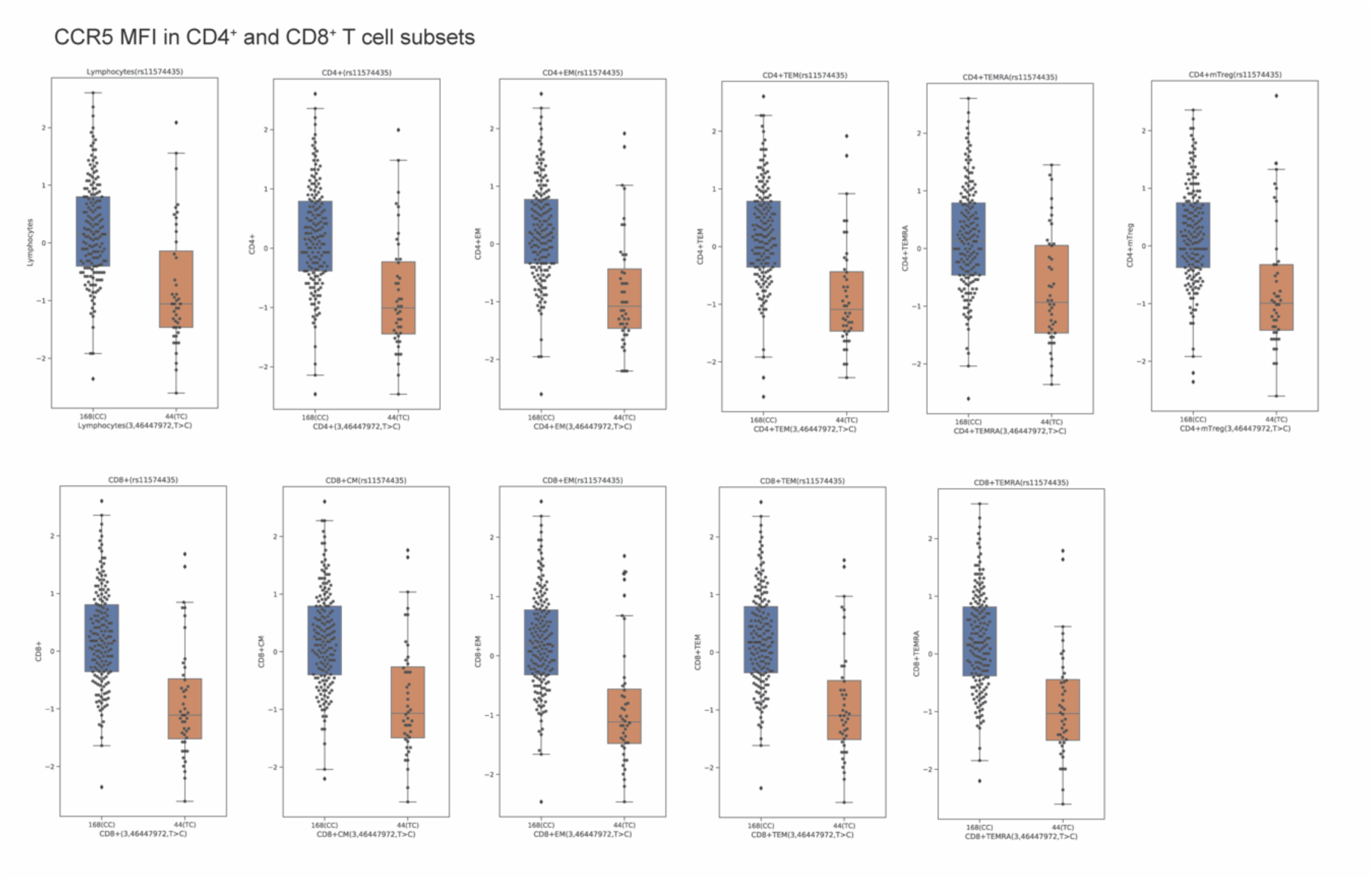
Genetic effect of rs11574435 SNP on CCR5 MFI in subpopulations of CD4^+^ and CD8^+^ T cells of PLHIV. Boxplots of CCR5 MFI levels in the different subsets of CD4^+^ and CD8^+^ T cells stratified according to the rs11574435 genotypes in PLHIV (P-value = < 0.05). Horizontal line in the boxplot, median; ends of the boxes, upper and lower quartiles.

Confounding these associations, rs11574435 SNP were shown to be highly in LD with *CCR5d32* (rs333) (R^2^ = 0.8423, D’ = 0.9591, P-value < 0.0001, European cohort of the 1000 Genome Project). After performing a Fisher’s exact test in order to study which of the alleles of the rs11574435 SNP was linked with the *CCR5d32*, we identified that the rs11574435-T allele was significantly linked with the presence of *CCR5d32* (odds ratio = 0.004 and P-value < 2 × 10^-16^). In addition, the rs71327064 SNP, associated with CD4^+^ TEMRA cells was also moderately linked to *CCR5d32* (R^2^ = 0.3379, D’ = 0.9062, P-value < 0.0001) (S1 Table).

The *CCR5d32* is a well-known causal variant affecting CCR5 expression. It is a structural variant that results in deletion of 32 base pairs of *CCR5* gene open reading frame and is associated with slower disease progression in PLHIV (5). The effects of this deletion in HIV susceptibility have been attributed to reduced expression of a functional CCR5 receptor (15). We therefore assessed the presence of *CCR5d32* in PLHIV and its effect on the CCR5 expression of the various cell subsets. A heterozygous (WT/delta32) phenotype for *CCR5d32* was found in 18,8% (*n=*40) of PLHIV, whereas 81,2% (*n=*173) did not have the deletion (WT/WT) (S5 Fig). With the exception of nTregs, all T cell subsets and monocytes from PLHIV carrying the WT/delta32, expressed significantly lower CCR5 MFI than those which are WT/WT PLHIV (Wilcoxon Test, P-value < 0.05) (S6 Fig). Moreover, the rs333 also influenced CCR5 proportions as subpopulations of CD4^+^ and CD8^+^ T cells of WT/delta32 PLHIV showed lower CCR5 proportions than WT/WT. No differences in CCR5 proportions were observed for nTregs and monocytes (S7 Fig).

**Fig 5.**
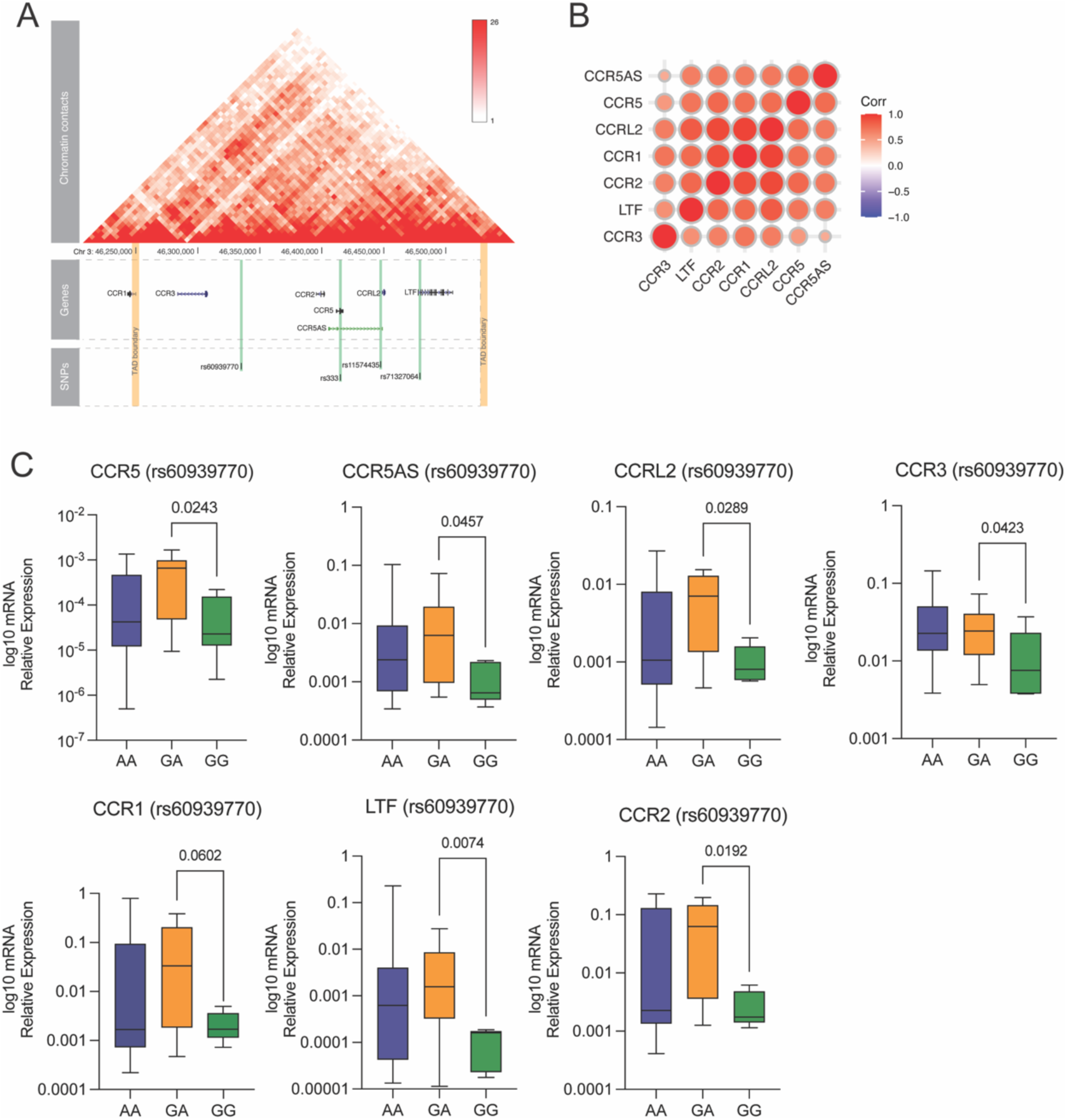
Chemokine receptors that are part of the *CCR5* gene cluster. (A) Topologically associating domain (TAD) of CCR5 locus. (B) Spearman’s correlation as the measure of similarities between the pattern mRNA expression of *CCR1, CCR3, CCR2, CCRL2, LTF, CCR5* and *CCR5AS* assessed by RT-PCR. Red indicates a strong positive correlation, whereas blue indicates a strong negative correlation (n= 58 PLHIV). (C) mRNA levels of *CCR1, CCR3, CCR2, CCRL2, LTF, CCR5* and *CCR5AS* were determined by RT-PCR and the values were stratified based on rs60939770 genotypes. Data were analysed using Mann-Whitney U-test (P-value < 0.05). Horizontal line in the boxplot, median; ends of the boxes, upper and lower quartiles.

**Figure 6.**
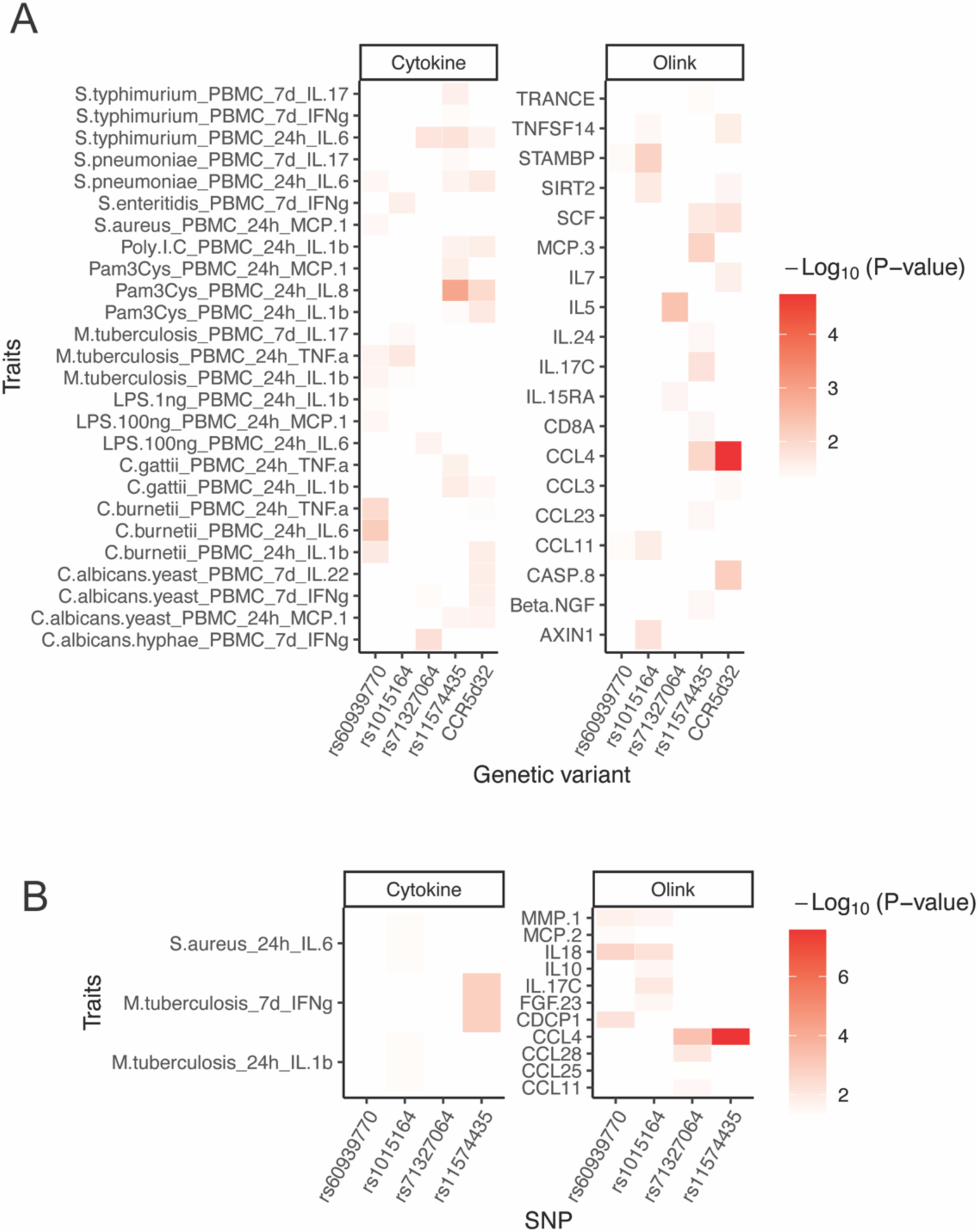
The association results of all genome-wide significant QTLs and *CCR5delta32* with inflammatory markers. (A) Correlation results between the SNPs associated with CCR5 surface expression and cytokines produced by PBMCs and circulating proteins (Olink) in PLHIV and (B) healthy controls (300BCG healthy cohort). The color represents the correlation P-value transformed by -Log10. The correlations were estimated using a linear model with age (at visit) and sex as co-factors.

To evaluate whether the SNPs we identified (rs11574435 and rs71327064) were independently causal to decreased CCR5 MFI in PLHIV, we stratified the six individuals carrying only the rs11574435 SNP and not *CCR5d32.* We found no significant differences in CCR5 MFI expression on CD4^+^ T cells, nor in the majority of CD8^+^ T subsets and monocytes of PLHIV carriers of rs11574435-TC without *CCR5d32* (WT/WT) compared to those carrying neither rs11574435 nor *CCR5d32.* CD8^+^ effector memory T cells was the exception, which a significant increase in CCR5 MFI expression of PLHIV carrying the rs11574435-TC was observed in comparison to the individuals rs11574435-CC and *CCR5d32* (WT/WT) (S8 Fig).We applied the same stratification strategy to evaluate the effects of CCR5 MFI in CD4^+^ TEMRA cells of individuals carrying the rs71327064 SNP only in comparison to individuals with neither rs71327064 SNP nor *CCR5d32.* We observed that in the absence of *CCR5d32,* CCR5 MFI expression was significantly lower in individuals carrying the rs71327064 SNP only than in individuals with neither rs71327064 nor *CCR5d32* (Wilcoxon Test, P-value < 0.05) (S9 Fig). Together, these results suggest that *CCR5d32* is playing the major effect in CCR5 MFI expression in relation to rs11574435 in the majority of on CD4^+^ and CD8^+^ T subsets evaluated, whereas the rs71327064 may modulate CCR5 MFI in CD4^+^ TEMRA cells independently of the presence of *CCR5d32*. In addition, PLHIV carrying the combination of the rs11574435 and rs71327064 SNPs together with the *CCR5d32* (WT/delta32) had a remarkable decrease in CCR5 MFI expression in comparison to the individuals without any of the three genetic variants (Wilcoxon Test, P-value < 0.05). These findings shows that the identified SNPs exert their functions in a cell-type specific manner, a feature that was only possible to explore due the detailed assessment of CCR5 expression on the surface of different subsets of T cells.

### 2.5 rs60939770 influence the mRNA expression of other nearby chemokines receptors within the *CCR5* gene cluster

We performed *cis*-expression QTL (eQTL) mapping using RNA sequencing data from whole blood samples of healthy individuals (16). The results showed that the genetic variants we identified in the *cis-*region to CCR5 locus is not only associated with the expression of *CCR5* but also influences the expression of other nearby chemokines genes (Fig. 2, S1 Table). Transcription of coregulated genes occurs in the context of long-range chromatin interactions, which genes and transcriptional regulatory elements, such as long noncoding RNAs are brought into close proximity to regulate protein-coding gene expression (17). We sought to map the three-dimensional chromatin structures, known as topologically associating domain (TAD) of CCR5 locus, and test which genes within this genomic neighbourhood may be subjected to such long-range co-regulatory mechanisms and investigate the correlations of long-range contact with transcription in PLHIV. Publicly available genome-wide chromosome conformation capture (Hi-C) data obtained from K562 cells revealed *CCR1, CCR3, CCR2, CCRL2, LTF,* and *CCR5AS* located within the same TAD as *CCR5* (Fig. 5A). In order to understand the co-regulatory relationship between these genes at the transcriptional level, we next profiled the expression of *CCR1, CCR3, CCR2, CCRL2, LTF, CCR5,* and *CCR5AS* transcripts by qPCR on whole blood samples of individuals from PLHIV. At baseline, without the presence of any stimulants, we observed a similar pattern of expression in between the *CCR5* and the nearby genes identified in the TAD (Fig. 5B, Spearman’s correlation, P-value < 0.05).

As the eQTL analysis of candidate genes performed in whole blood of healthy individuals revealed that the rs60939770 is associated with the genes located within the CCR5 TAD (Fig. 2), we next assessed the effects of rs60939770 in modulating the expression of *CCR1, CCR3, CCR2, CCRL2, LTF, CCR5,* as well as *CCR5AS* in whole blood of PLHIV. PLHIV carrying the rs60939770-GG genotype had lower expression of *CCR1, CCR3, CCR2, CCRL2, LTF, CCR5* mRNA and *CCR5AS* RNA in comparsion to the individuals rs60939770-GA (Wilcoxon Test, P-value < 0.05 (Fig. 5C). We also evaluated the effects of rs11574435 and in modulating the expression of *CCR5AS, CCR5* and nearby genes in PLHIV carrying the different SNP genotypes. In subjects with the rs11574435-TC genotype, a significant increase (Wilcoxon Test, P-value < 0.05) in *CCR5, CCR3, CCRL2* and *LTF* mRNA expression in comparison to rs11574435-CC. Although no significant differences were observed for *CCR5AS*, *CCR1* and *CCR2,* there was a tendency to higher expression in TC genotypes (S10 Fig). Together, these findings suggest that the *cis*-genetic regulators of CCR5 that lead to the modulation of CCR5 surface expression (measured as cell proportions or MFI) also modulate mRNA expression of *CCR5* and other chemokines receptors within the same locus.

### 2.6 The genetic regulation of CCR5 expression influences the cytokine production capacity of circulating immune cells

CCR5 expression is known to facilitate the chemotaxis of immune cells to sites of infection or inflammation, a process that results in amplification of the inflammatory responses (18). We therefore tested whether the SNPs that are associated with CCR5 surface expression (rs60939770, rs1015164, rs71327064, rs11574435 and *CCR5delta32*) were related with the production capacity of inflammatory cytokines and chemokines after ex vivo stimulation of PBMCs in our 200HIV cohort (19). We found that the rs60939770 is associated with the production of the innate immune cells-derived soluble mediators, IL-1β, IL6, TNF and monocyte chemoattractant protein-1 (MCP-1) in PLHIV (nominal P-value < 0.05). Of note, despite the fact of rs1015164 being in LD with rs60939770, we demonstrated that in terms of functions these two SNPs differ, as rs1015164 was associated with the production of the TNF and IFNγ (Fig. 6A). When we further assessed the SNPs identified to be associated with CCR5 MFI levels, we observed that the rs71327064 was associated with the production of both IL-6 and IFNγ. Moreover, the rs11574435 SNP was significantly associated with the production of both innate and adaptive inflammatory mediators of PBMCs of PLHIV, including MCP-1, IL-1β, TNF, IL8, IL6 and IFNγ, IL17, respectively (Fig. 6A). In the HC cohort, rs11574435 SNP was also associated with the levels of IFNγ by PBMCs (Fig. 6B). Of importance, *CCR5delta32* was also shown to modulate the production of MCP-1, IL-1β, IL8, IL6 and IFNγ and IL-22.

Next, we tested the association between the identified SNPs and the concentrations of circulating inflammatory mediators both in PLHIV and in HC. We observed that the rs11574435 is associated with the secretion of beta-chemokines including the C-C Motif Chemokine Ligand 4 (CCL4), which together with CCL3 and CCL5 are known as anti-CCR5 factors (20). These chemokines downregulate or block the receptors CCR5, CCR1 and CCR2 resulting in HIV infection inhibition (21). Besides CCL4, rs11574435 SNP was associated with the levels of the chemokines, MCP-3, CCL23, CCL11 as well as the production of SCF, IL17C, CD8A in PLHIV (Fig. 6A). *CCR5delta32* was also related to the production of CCL4 and SCF, but not with MCP-3, CCL23, CCL11 or IL17C, CD8A. In addition, *CCR5delta32* was associated with the production of caspase 8. In the HC cohort, rs11574435 SNP is also associated with circulating CCL4 concentrations (Fig. 6B). Thus, despite the co-occurrence of *CCR5delta32* and rs11574435 as the main genetic factors responsible for the modulation of CCR5 MFI expression, these genetic variants alter the production of soluble mediators in a different manner. Our findings indicate that the differences in CCR5 expression is associated with the levels of immune activation in PLHIV, which may have an impact on the pathogenesis of HIV infection.

### 3. Discussion

In the present study, we identified three novel common genetic loci that were associated with cell-type-dependent surface expression of CCR5 in two independent cohorts from Western European descent, one consisting of virally suppressed PLHIV, the other cohort including healthy individuals. We also show that these genetic variants not only affect CCR5 expression but also other genes that are part of the same topologically associating domain in the CCR5 locus. Finally, our results indicate that these genetic variants cause altered inflammatory responses.

The strongest genome-wide significant locus was the rs60939770 variant in the *cis*-region of CCR5 locus, which was found to be specifically associated with the percentage of CCR5^+^CD4^+^ mTregs, both in PLHIV and in healthy controls. Tregs cells are important to control HIV replication by reducing T-cell activation, which decreases the availability of target cells for HIV (22). Moreover, in agreement with our findings, previous studies have shown that memory T cells express higher CCR5 in comparison to naive T cells (23). Memory Tregs may alleviate tissue damage during pro-inflammatory conditions and CCR5 expression on these cells may direct them into inflamed tissues (24). The precise role of Tregs in the HIV infections remains an extensive topic of discussion. The rs60939770 SNP is not correlated with the well-described *CCR5delta32* (rs333), but with another common variant, rs1015164, also in the *cis*-region of CCR5 locus. The latter variant is linked with HIV progression parameters such as viral load and CD4+ T cell counts (12, 25). Our data indicate that CCR5 expression of mTregs is also affected by rs1015164, therefore further investigation will be needed to determine which of the two genetic variants from this locus is causal for CCR5 modulation.

Apart from analyzing whether or not immune cells express CCR5, we also assessed the intensity of CCR5 surface expression. Here we found two different genetic variants, rs11574435 and rs71327064, associated with the MFI of surface CCR5 molecules in subpopulations of CD4^+^, CD8^+^ T cells and CD4^+^ TEMRA cells, respectively. rs11574435 is in LD with *CCR5delta32* and it is located in a transcript called *CCR5AS.* CCR5 expression is low in naïve cells, but with cell differentiation as well as stimulation, the surface expression of CCR5 on immune cells increases (23, 26). This is in line with our data which shows that the rs11574435 SNP and *CCR5delta32* variants associate with CCR5 MFI expression in mostly differentiated CD4^+^ and CD8^+^ T cells. Interestingly, unlike rs11574435, the *CCR5delta32* affected CCR5 MFI expression also in naive T cells as well as monocytes. Epigenetic factors may play a role in this, as the DNA methylation content of *CCR5* is different in naive vs differentiated cells (10). Of note, among PLHIV we observed that not all subjects carry both rs11574435 and *CCR5delta32* and we were able to study their effects separately. Our results indicate that rs11574435 has no influence in CCR5 surface expression, which we have shown to be attributed to the effects of *CCR5delta32*. This differed when we looked into the effects of rs71327064, which was identified associated with CCR5 MFI on CD4^+^ TEMRA. The presence of rs71327064 in CD4^+^TEMRA led to decreased CCR5 MFI surface expression. CD4+TEMRA cells are differentiated effector memory CD4 cells that highly express CCR5, correlate with CD4^+^ T cells numbers but are resistant to R5-tropic HIV-1 (27). CD4^+^ TEMRA may therefore be a resistant subset of T cells to HIV infection, which might have a beneficial role during HIV infections.

One may question whether CCR5 expression intensity is as relevant for HIV susceptibility, compared to the presence or absence of CCR5 on the cell surface. CD4 and CCR5 should co-localize so HIV can infect the cell. It has been shown, however, that CCR5, CXCR4 and CD4 are predominantly present on microvilli in different cell types, including T cells and macrophages and that these microclusters of CD4 and chemokine receptors were frequently separated by less distance than the diameter of an HIV virion (28), indicating that cells with low CCR5 expression may still be susceptible for HIV infection. Although our findings concerning the genetic modulation of CCR5 expression in the different cell substes are new, we have not investigated whether this would also be relevant for the stablishement or susceptibility of HIV infections. Moreover, long-lasting subsets of T cells, including memory T cells expressing CCR5, are known to host HIV during latency, named also as viral reservoirs (29). Therefore, the genetic variants we identified associated to T cells with memory functions might have important implication in the development of persisting viral reservoirs.

Memory T cells are known to be the main responders to beta-chemokines, and high expression of CCR5 on quiescent cells prompt them to be highly responsive to chemokine gradients at sites of immune and inflammatory responses (30, 31). Therefore, cells expressing higher levels of CCR5 can amplify inflammation favoring the development of non-AIDS comorbidities such as cardiovascular diseases (32). Importantly, we have demonstrated that rs60939770 and rs11574435 were not only associated with the proportions of CCR5 positive cells and MFI on the surface of memory CD4^+^ and CD8^+^ T cells, but also with *CCR5* mRNA levels and the expression of other nearby chemokines receptors (CCR1, CCR2, CCR3) which are part of the same TAD. Of note, similar to CCR5, CCR2 has been described to enhance HIV infection (33). As the rs60939770 and rs11574435 SNPs are located within non-coding regions of the genome, it is likely that all chemokines within the same TAD share a common regulator which influences their expression in a similar manner. Also, as these regulators are stimulus and cell-type specific (34), it is worthy considering that the transitional state of naïve T to mature memory cells trigger the induction of such elements leading to the modulation of CCR5 expression. We have shown that genetic variants might play a role in the expression of these regulators, however further studies are required to explore the relationship of these non-coding transcripts and the expression of CCR5 and other chemokine receptors.

Similar effects observed for rs11574435 were seen for the individuals carrying the *CCR5delta32*. The *CCR5delta32* has been previously associated with differential expression of chemokine receptors coding genes, for example *CXCR2, CCRL2,* as well as genes involved in T cell activation (CD6) and maturation (CD7) (35). In addition, we have demonstrated that rs11574435 as well as *CCR5delta32* are related to the production of other soluble factors including chemokines and inflammatory cytokines. Previous studies have indeed shown that CCR5 is a cell surface signaling receptor that plays a role in activation of inflammatory genes (36). The chemokines CCL3, CCL4, and CCL5/RANTES, known as CCR5 ligands, may protect CD4 T cells from HIV infection (20). Of note, CCL5 and CCL4 also bind to different chemokine receptors (37). Here we observed that rs11574435 and *CCR5delta32* modulate the production of CCL4 in PLHIV. Moreover, decreased CCR5 surface expression triggered the increased *CCR5* mRNA expression likely due to feedback mechanisms. Altogether, our data suggest that certain genetic variants do not only affect CCR5 expression levels, but also production of chemokines, which may directly impact viral entry or modulate immune responses associated with HIV pathogenesis including non-AIDS comorbidities.

The current study has several limitations. First, despite large sample sizes in two cohorts, there is a limited power in detection of small genetic effects. Second, a single monoclonal anti-CCR5 antibody (2D7) was used to detect CCR5 and certain isoforms and CCR5 conformations may have not been recognized by this antibody (38). However, compared to other antibodies, 2D7 probably reacts best with conformations of CCR5 that are relevant to HIV-1 entry (39). Third, we did not evaluate the transcription factor FoxP3 to phenotype regulatory T cells. On the other hand, we have systematically compared the individuals from both PLHIV and HC cohorts in the same manner using other additional marks including CD25 and CD45RA for the assessment of naive and memory status (40). In addition, we have shown comparable results in between Tregs indentifed using FoxP3^+^Helios^+^ and CD25 (19).

In summary, the results presented herein indicate that genetic factors contribute to the interindividual variability of CCR5 surface expression in different subsets of immune cells of peripheral blood of both PLHIV and healthy controls of European ancestry. Furthermore, we show that the expression of certain chemokines (CCL4) chemokines receptors (*CCR1*, *CCR2*, *CCR3, CCRL2*), which are part of the same topologically associating domain, are also affected by these genetic factors.

## 4. Methods

### 4.1 Ethics

The study protocols of the 200 HIV pilot study and 300BCG were approved by the Medical Research Ethical Committee Oost-Nederland (ref. 42561.091.122 and NL58553.091.16, respectively) and conducted in accordance with the principles of the Declaration of Helsinki. All study participants provided written informed consent.

### 4.2 Study population

The volunteers of this study are part of the HFGP (www.humanfunctionalgenomics.org) (33). Between 14 December 2015 and 6 February 2017, individuals living with HIV were recruited from the HIV clinic of Radboud university medical center, the Netherlands. Inclusion criteria were Dutch/Western-European ethnicity, age ≥ 18 years, receiving cART > 6 months, and latest HIV-RNA levels ≤200 copies/ml. Exclusion criteria were: signs of acute or opportunistic infections, antibiotic use <1 month prior to study visit, and active hepatitis B/C. General baseline characteristics of PLHIV including CD4 Nadir and HIV RNA Zenith were described previously (19). The healthy individuals were included in the 300BCG study between April 2017 and June 2018 in the Radboud university medical center, the Netherlands. Exclusion criteria were use of systemic medication other than oral contraceptives or acetaminophen, use of antibiotics 3 months before inclusion, and any febrile illness 4 weeks before participation (13).

### 4.3 Genotypes, imputation, and quality control

For the PLHIV cohort, the genotyping, imputation, and quality control were described previously (41). In addition, for the healthy individuals, the genotyping, imputation, and quality control were performed as in the previous study (42).

### 4.4 QTL mapping

Firstly, the immune phenotypes (cell proportions and CCR5 levels) were transformed using inverse-normal transformation. Then, we used the R/MatrixEQTL package (43) to conduct the QTL mapping for the geometric mean of fluorescence intensity of CCR5 protein and CCR5 positive cell proportion in PLHIV and HC, respectively. We used a linear regression model with age and sex as co-variables. To evaluate the inflation of the summary statistics, we calculate the genomic inflation factor for each association analysis: lambda values vary between 0.980 and 1.017 (median = 1.0045, mean = 1.0018, stdev = 0.0069). Finally, we used P-value < 5 × 10^-8^ as the genome-wide significant threshold to select SNPs for downstream analysis. To visualize the identified association signals, we used circus (44) (v 0.69, Perl 5.028001) package to show the Manhattan plot of analysis results along with the genome coordination only for chromosome 3. The hierarchical tree from the clustering analysis of immune cell proportions is plotted using the R/ggtree (45) (v1.8.2) package powered by BioConductor (v3.10). To investigate the genomic context around genome-wide significant associations, summary statistics for each phenotype were uploaded to the LocusZoom (46) server to visualize regional QTL mapping scan results, using hg19 and 1000 Genomes Nov 2014 EUR as reference genome and for LD calculation, respectively. Manhattan plot and Q-Q plot for each association analysis were generated by package qqman using default parameter settings. Analysis using MatrixEQTL and qqman were performed using R language (v3.6.0) and the in-house scripts for preprocessing using Python (v3.7.0) language are hosted on GitHub https://github.com/zhenhua-zhang/qtl_mapping_pipeline.

### 4.5 Antibodies and flow cytometry

Flow cytometry analyses were conducted to assess CCR5 expression levels on several subsets of circulating immune cells (S1 Fig). Pre-processing stages including cell-processing and staining were similarly performed and by the same personnel in PLHIV and healthy controls. Venous blood was collected in sterile EDTA tubes. Details regarding cell processing and staining were described previously (47).

A Sysmex XN-450 automated hematology analyzer (Sysmex Corporation, Kobe, Japan) was used for determination of cell counts and to calculate absolute numbers of CD45+ white blood cell (WBC) counts as measured by flow cytometry. Methods, S3 Table shows the fluorochrome conjugates and clone identity of the antibodies. Flow cytometry data were acquired using a 10-color Navios flow cytometer (Beckman Coulter) and the Kaluza Flow Cytometry software (Beckman Coulter, version 2.1). Different subsets of immune cells were identified by sequential manual gating (S11 Fig).

### 4.6 Molecular genotyping of *CCR5d32*

Whole blood samples of PLHIV of the 200 HIV pilot study collected in EDTA tubes (BD Vacutainer) were used for genomic DNA extraction. The assessment of the region of the CCR5 gene containing the d32 deletion was adapted from (48). Primer sequences are listed in Methods, S4 Table. The PCR reactions were prepared using the 5X Q5 buffer, 10 mM dNTPs, Q5 High-Fidelity DNA Polymerase (New England Biolabs, Inc) and 10 uM forward and reverse primers. 50 ng of DNA was used as template. The PCR protocol consisted of 1 cycle of 98℃ for 30s, 35 cycles of 98℃ for 10s, 62℃ -10s, 72℃ -10s and 1 cycle of 72℃ for 2min. Fragments obtained from PCR were separated in 2% agarose gel containing ethidium bromide for visualization.

### 4.7 RNA isolation and quantitative real time-PCR

RNA was extracted from the whole blood of PLHIV of the 200 HIV pilot study using the QIAGEN PAXgene Blood RNA extraction kit (QIAGEN, Netherlands) according to the instructions of the manufacturer. Subsequently, RNA was reversely transcribed into cDNA by using iScript (Bio-Rad, Hercules, CA, USA). Diluted cDNA was used for qPCR that was done by using the StepOnePlus sequence detection systems (Applied Biosystems, Foster City, CA, USA) with SYBR Green Mastermix (Applied Biosystems). The mRNA and RNA relative expression analysis was done with the 2^-dCt method and normalized against the housekeeping gene RPL37A. Primer sequences are listed in Methods, S4 Table.

### 4.8 PBMCs stimulation experiments and plasma proteomics

Venous blood was collected in EDTA tubes (BD Vacutainer) and PBMCs were obtained by density centrifugation over Ficoll-Paque (VWR, Amsterdam, the Netherlands). Freshly isolated PBMCs (0.5 million cells/well) were incubated with different stimuli including bacterial (*Staphylococcus aureus*, *M. tuberculosis*, *Streptococcus pneumoniae*, *Coxiella burnetii, Salmonella enteritidis, Salmonella typhimurium*), fungal (*Cryptococcus gattii, Candida albicans* hyphae and yeast) and other relevant antigens (Poly:IC (100 ug/mL - Invivogen; TLR3 ligand), *E. coli* LPS (1 and 100 ng/mL - Sigma; TLR4 ligand) and Pam3Cys, (10 ug/mL - EMC microcollections; TLR2 ligand)), in round-bottom 96-well plates (Greiner Bio-One, Frickenhausen, Germany) at 37°C and 5% CO2 in the presence of 10% human pooled serum for lymphocyte-derived cytokines assessment. The concentration of the mentioned bacterial and fungal stimuli are described previously (49). Supernatants were stored at - 20°C. Levels of the monocytes-derived cytokines (TNF, IL-1β, IL-6) as well as chemokines (MCP-1, IL-8) were measured in the supernatants after 24 hours incubation. Levels of lymphocyte-derived cytokines (IFNγ, IL-17) were determined after 7 days (PeliKine Compact or Duoset ELISA, R&D Systems). Baseline inflammatory plasma markers from both cohorts, 200HIV pilot study and 300BCG, were measured by targeted proteomics as applied by the commercially available Olink Proteomics AB (Uppsala Sweden) Inflammation Panel (92 inflammatory proteins), using a Proceek © Multiplex Proximity extension assay. Expression levels were calculated as described by Koeken et al 2020 (13).

## Acknowledgments

The authors thank all volunteers from the Radboud University Medical Centre (Radboudumc) for participation in the study. Z.Z. is supported by a joint fellowship from the University Medical Center Groningen and China Scholarship Council (CSC201706350277) and the Singh-Chhatwal-Postdoctoral Fellowship at the Helmholtz Centre for Infection Research. We thank the UMCG Genomics Coordination center, the UG Center for Information Technology, and their sponsors BBMRI-NL & TarGet for storage and compute infrastructure. Y.L. was supported by an ERC Starting Grant (948207) and the Radboud University Medical Centre Hypatia Grant (2018) for Scientific Research. MGN was supported by an ERC Advanced Grant (833247) and a Spinoza Grant of the Netherlands Organization for Scientific Research

## Supporting information

**S1 Table.**
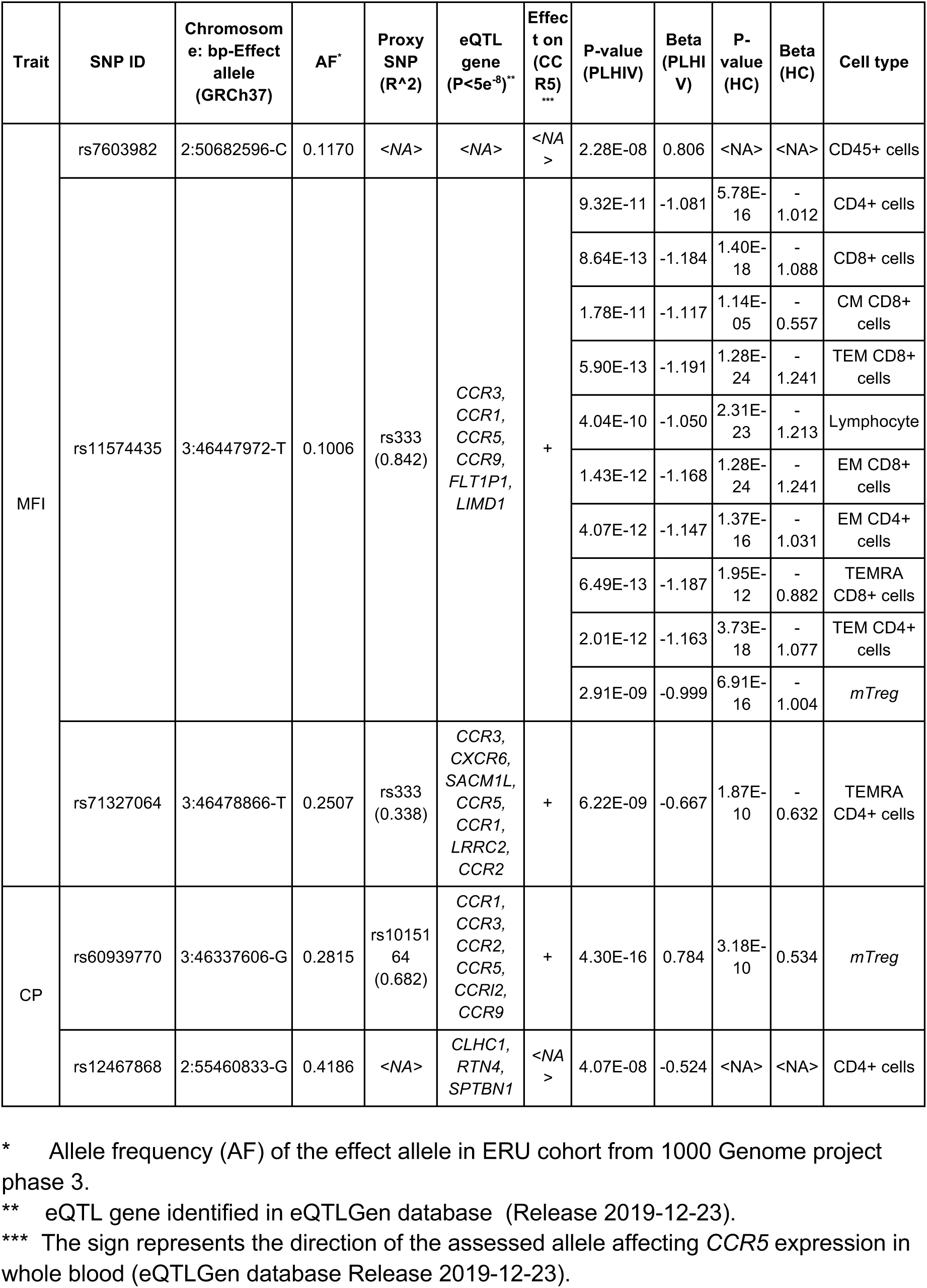
Genomic-wide significant CCR5 QTL SNPs in PLHIV. Abbreviations: CM = central memory, EM = effector memory cells (CD45RA-CCR7-), TEMRA = effector memory cells expressing CD45RA (CD45RA+CCR7-), and TEM = total effector memory (i.e. the total pool of effector memory cells).

**S2 Table.** Genomic-wide significant CCR5 QTL SNPs in healthy individuals. Abbreviations: CM = central memory, EM = effector memory cells (CD45RA-CCR7-), TEMRA = effector memory cells expressing CD45RA (CD45RA+CCR7-), and TEM = total effector memory (i.e. the total pool of effector memory cells).

| SNP ID | Effect allele | Trait | P-value | Beta | Cell type |
| --- | --- | --- | --- | --- | --- |
| rs113010081 | 3:46457412-C | MFI | 1.64E-15 | -1.039 | CD45+ cells |
|  |  |  | 6.96E-19 | -1.142 | CD4+ cells |
|  |  |  | 2.24E-27 | -1.355 | Lymphocytes |
|  |  |  | 9.47E-34 | -1.474 | EM CD8+ cells |
|  |  |  | 6.45E-20 | -1.171 | EM CD4+ cells |
|  |  |  | 1.19E-21 | -1.217 | TEM CD4+ cells |
|  |  | CP | 4.25E-13 | -0.949 | Lymphocytes |
|  |  |  | 4.85E-08 | -0.721 | TEMRA CD4+ cells |
| rs113341849 | 3:46384204-A | MFI | 2.07E-22 | -1.217 | CD8+ cells |
|  |  |  | 1.49E-29 | -1.376 | TEM CD8+ cells |
|  |  |  | 2.87E-14 | -0.969 | TEMRA CD8+ cells |
|  |  |  | 7.21E-15 | -1.001 | TEMRA CD4+ cells |
|  |  |  | 8.07E-18 | -1.088 | mTreg |
|  |  | CP | 2.68E-14 | -0.965 | CD4+ cells |
|  |  |  | 2.42E-13 | -0.907 | CD8+ cells |
|  |  |  | 6.30E-16 | -1.036 | TEM CD8+ cells |
|  |  |  | 1.41E-15 | -1.024 | EM CD8+ cells |
|  |  |  | 3.82E-22 | -1.202 | EM CD4+ cells |
|  |  |  | 1.55E-18 | -1.087 | TEMRA CD8+ cells |
|  |  |  | 5.45E-22 | -1.196 | TEM CD4+ cells |
|  |  |  | 7.36E-20 | -1.149 | mTreg |
| rs9670662 | 13:111098701-A |  | 2.60E-08 | 0.429 | CM CD4+ cells |

**S1 Fig.**
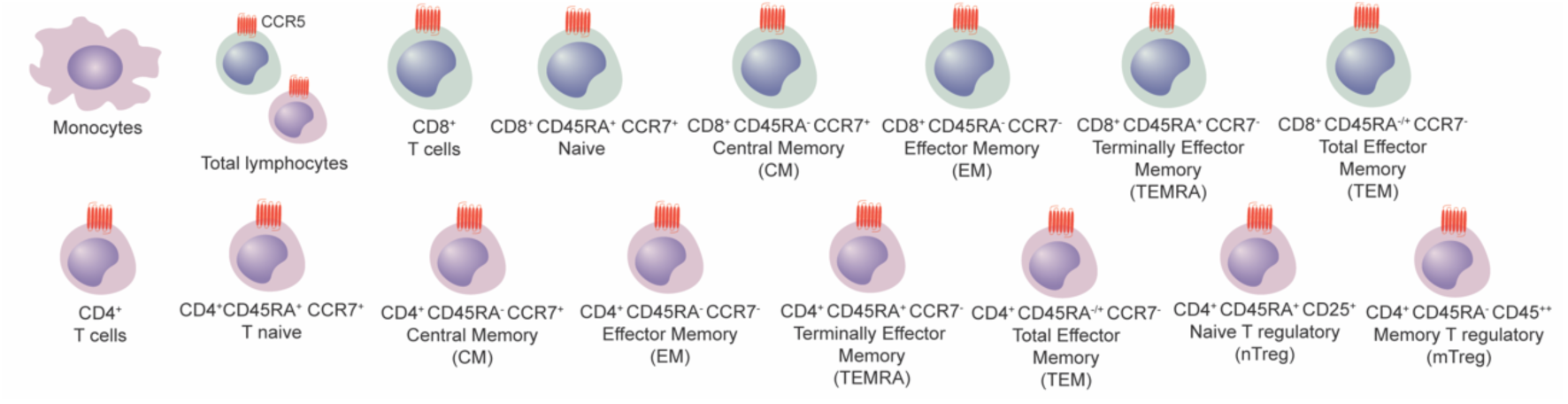
Circulating immune cells subpopulations in which CCR5 surface expression was assessed by flow cytometry.

**S2 Fig.**
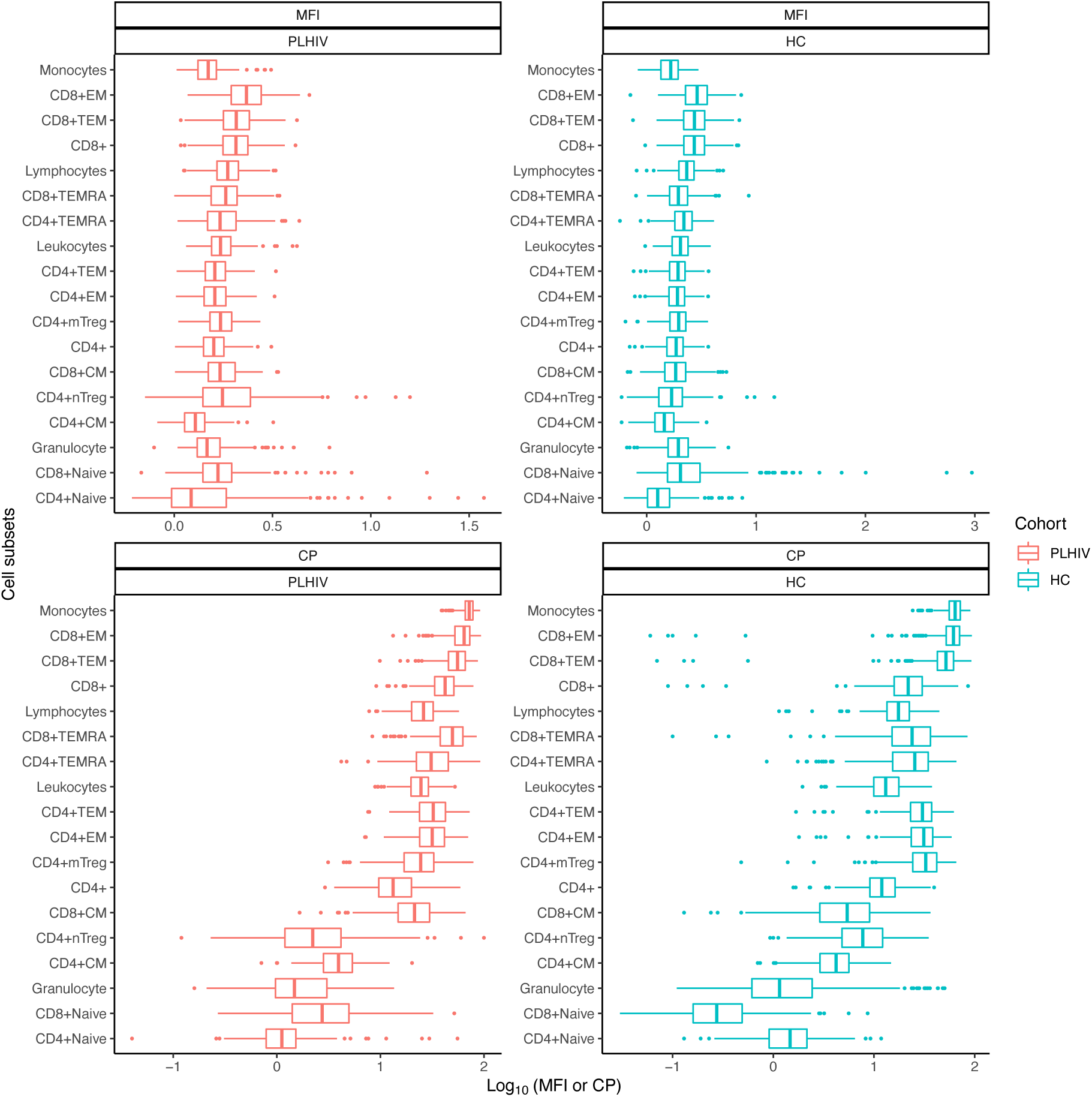
The distribution of CCR5 MFI and cell proportions (CP) measured in PLHIV (200HIV) and HC (300BCG).

**S3 Fig.**
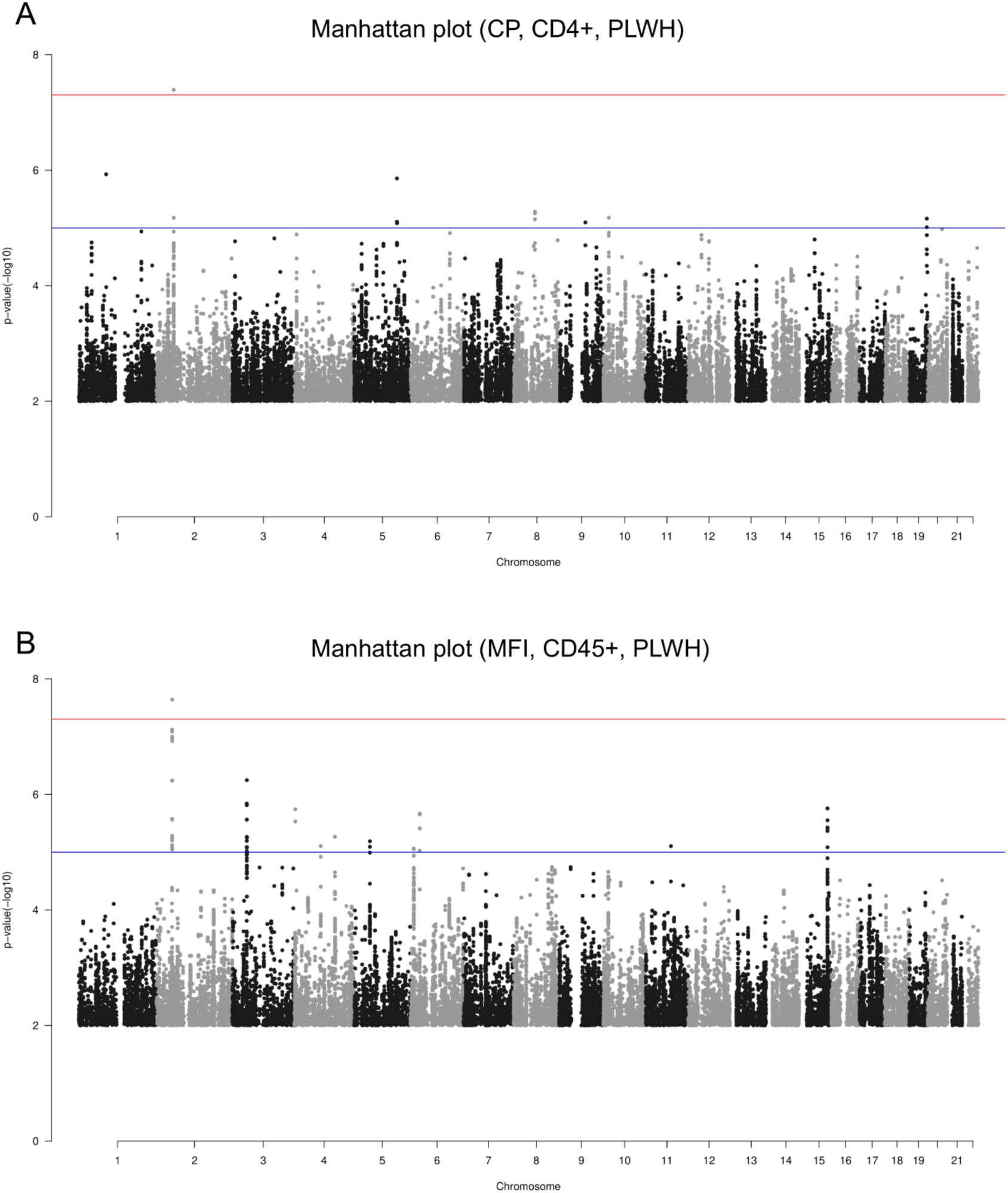
Manhattan plots shows two loci identified for CCR5 cell proportions in CD4^+^ cells (A) and for CCR5 MFI in CD45^+^ cells (B). Red lines correspond to a genome-wide significant threshold, whereas blue lines a represent suggestive threshold. Genomic variants are shown on the x-axis and y axis indicate the association between each variant and CD4^+^ cells (A) and MFI in CD45^+^ cells (B), respectively.

**S4 Fig.**
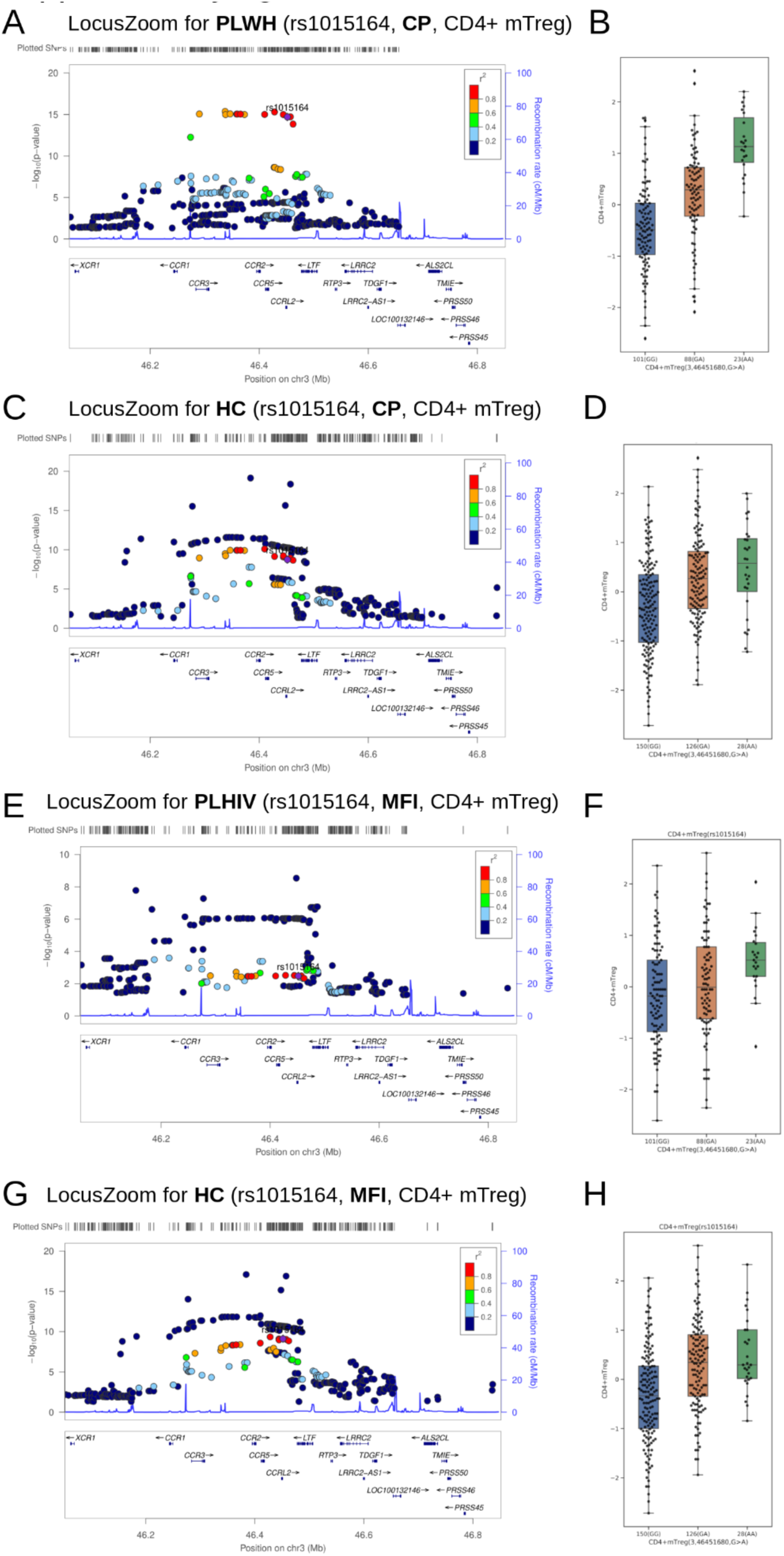
rs1015164 is associated with CCR5 cell proportions (CP) and MFI in mTreg cells from PLHIV and HC. (A) and (C) are regional plots (LocusZoom) showing rs1015164 associated with CP in CD4^+^ mTreg of PLHIV and HC, respectively. (B) and (D) are boxplots of CP in CD4^+^ mTregs stratified according to the genotypes of rs1015164 in PLHIV and HC, respectively. (E) and (G) are regional plots (LocusZoom) showing rs1015164 associated with MFI in CD4^+^ mTreg of PLHIV and HC, respectively. (F) and (H) are boxplots of MFI in CD4^+^ mTregs stratified according to the genotypes of rs1015164 in PLHIV and HC, respectively.

**S5 Fig.**
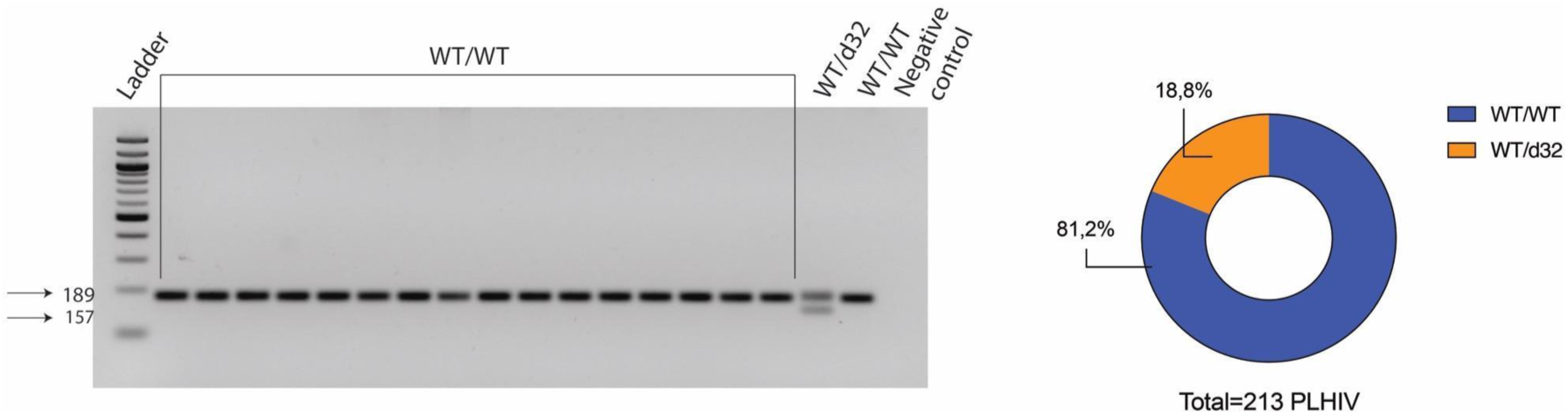
Schematic representation of how *CCR5d32* were assessed in individuals part of the 200HIV pilot study. WT allele is expected at 189bp and the *d32* is expected at 157bp. After the ladder in lane 1, lanes 2-16 represents homozygous wild-type genotype (fragment of 189bp), lane 17 represents a heterozygous genotype (fragments of 189bp and 157bp), lane 18 represents homozygous wild-type genotype and lane 19 is the negative control for PCR reaction. Part of a whole plot showing the distribution of WT/WT and WT/d32 in the entire cohort of HIV patients.

**S6 Fig.**
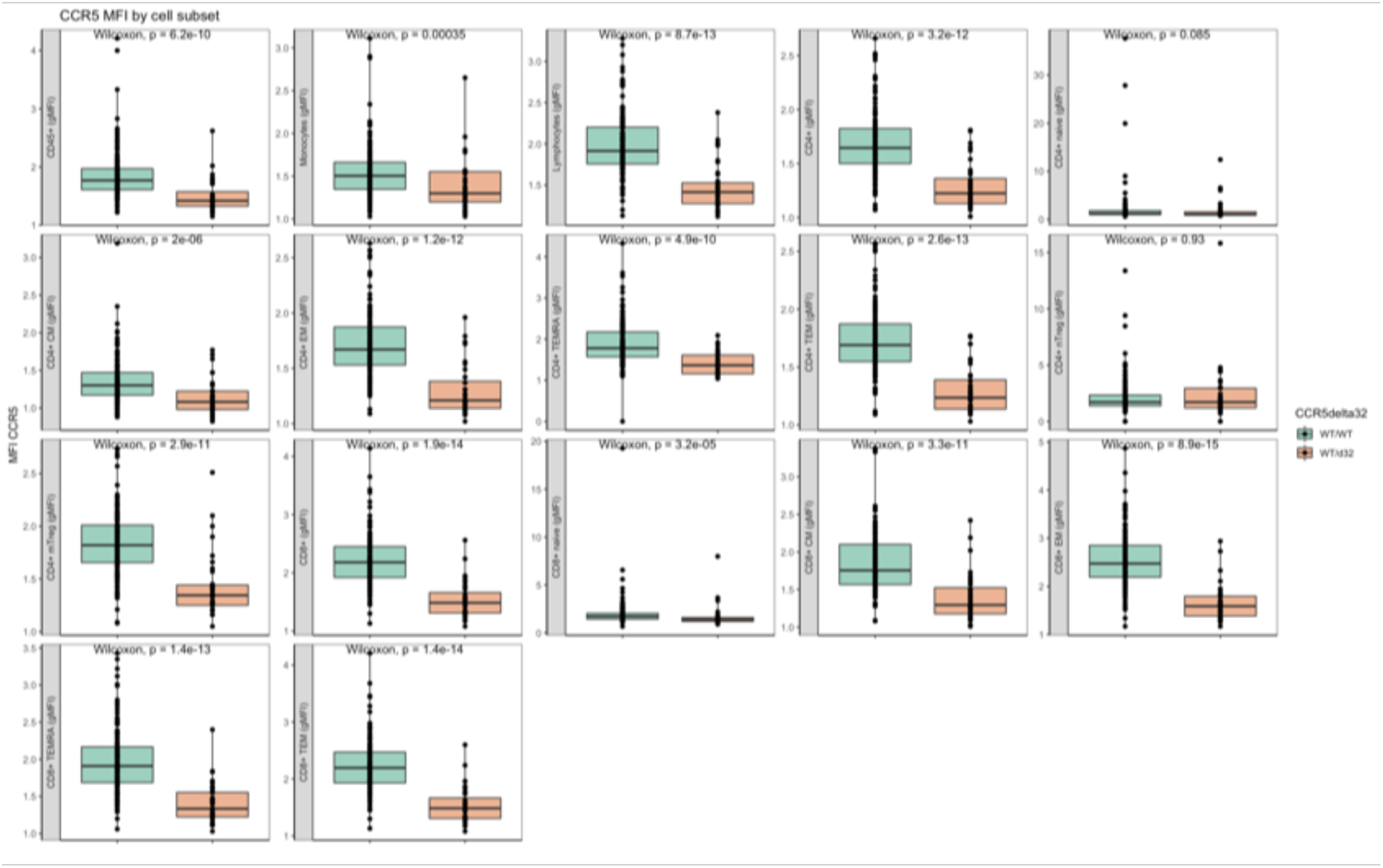
CCR5 geometric mean of fluorescence intensity (MFI) stratified based on *CCR5 delta32* (WT/WT= green, WT/d32= orange; all PLHIV). Data were analysed using Wilcoxon matched pairs signed-rank test (P-value < 0.05).

**S7 Fig.**
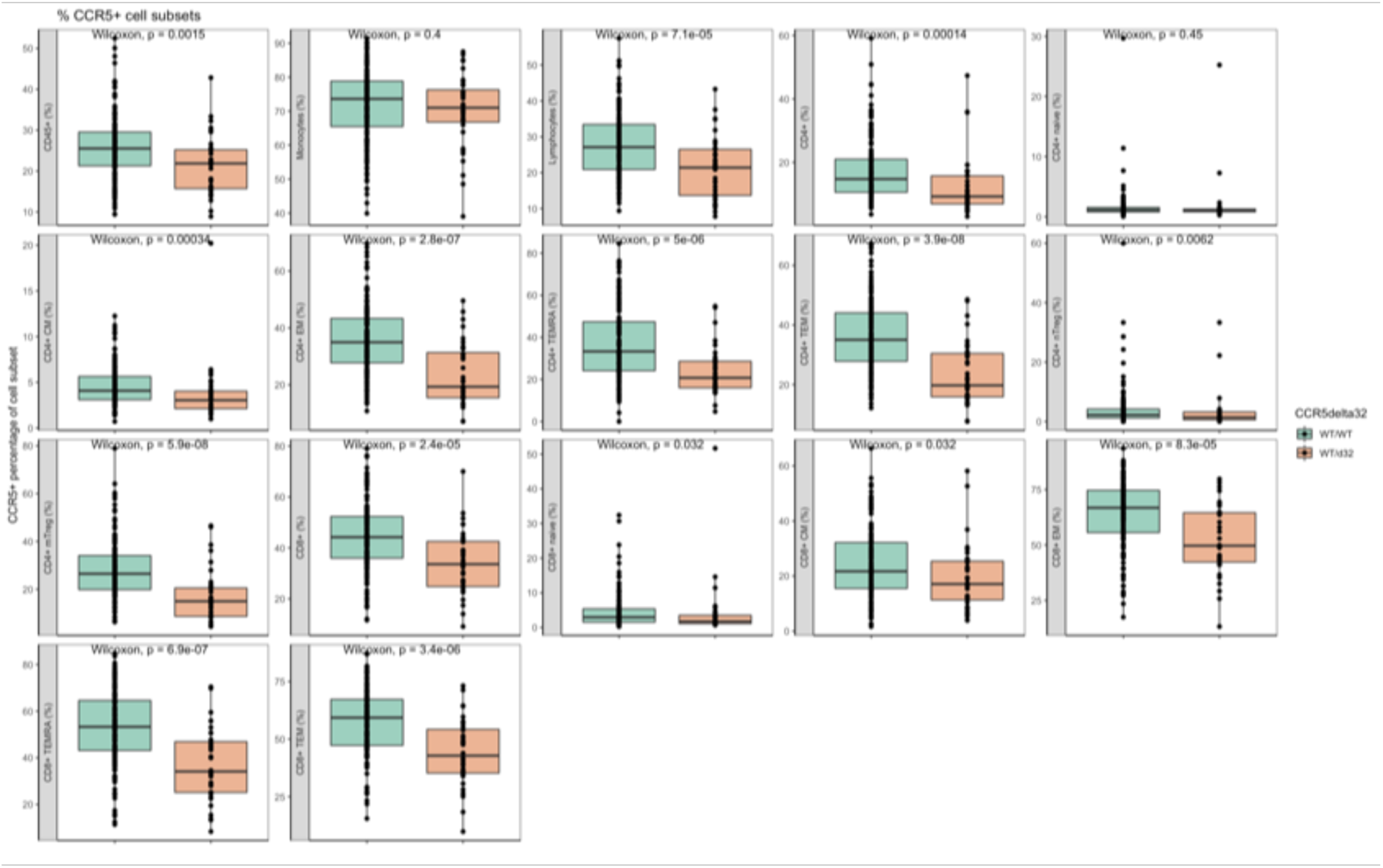
Percentages of CCR5 positive cells (cell proportions) stratified based on *CCR5 delta32* (WT/WT= green, WT/d32= orange; all PLHIV). Data were analysed using Wilcoxon matched pairs signed-rank test (P-value < 0.05).

**S8 Fig.**
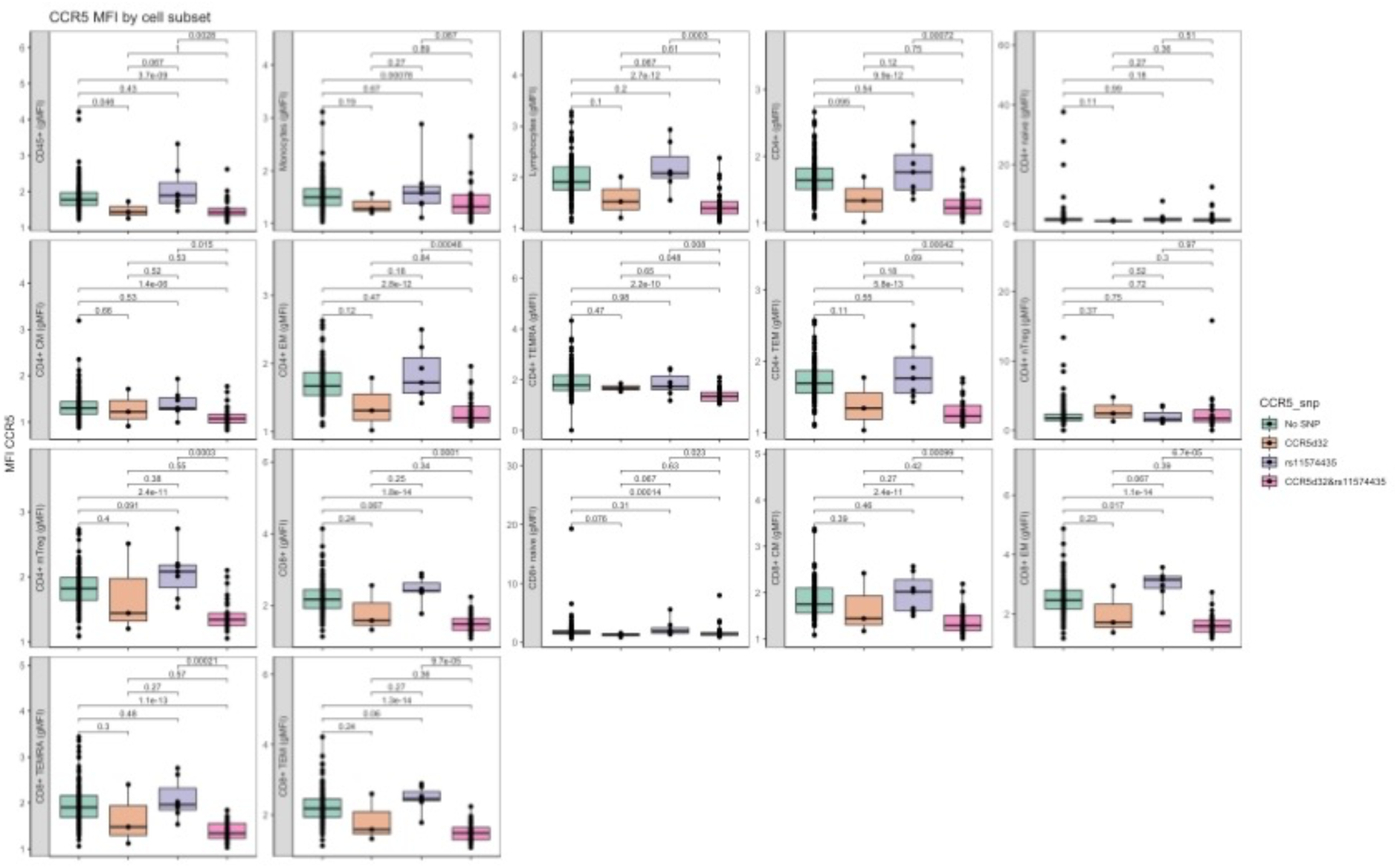
CCR5 MFI stratified based on individuals carrying no SNP (green), *CCR5 delta32* (orange) or rs11574435 (purple) only and both *CCR5 delta32*/rs11574435 together (pink). Data referred to PLHIV analysed using Wilcoxon matched pairs signed-rank test (P-value < 0.05).

**S9 Fig.**
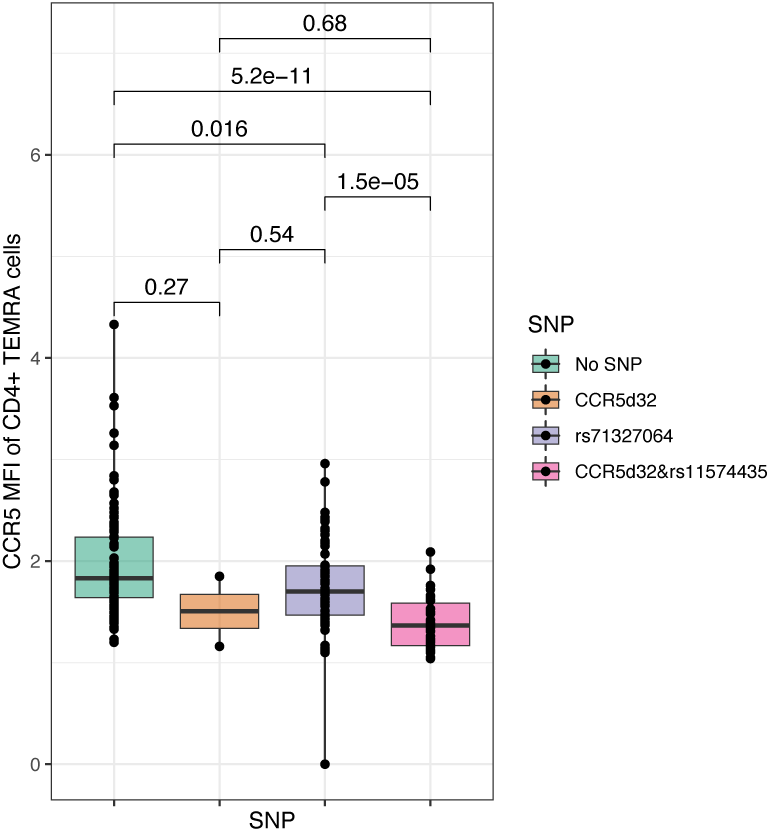
CCR5 MFI in CD4^+^ TEMRA cells stratified based on individuals carrying no SNP (green), *CCR5 delta32* (orange) or rs71327064 (purple) only and both *CCR5 delta32*/rs71327064 together (pink). Data referred to PLHIV analysed using Wilcoxon matched pairs signed-rank test (P-value < 0.05).

**S10 Fig.**
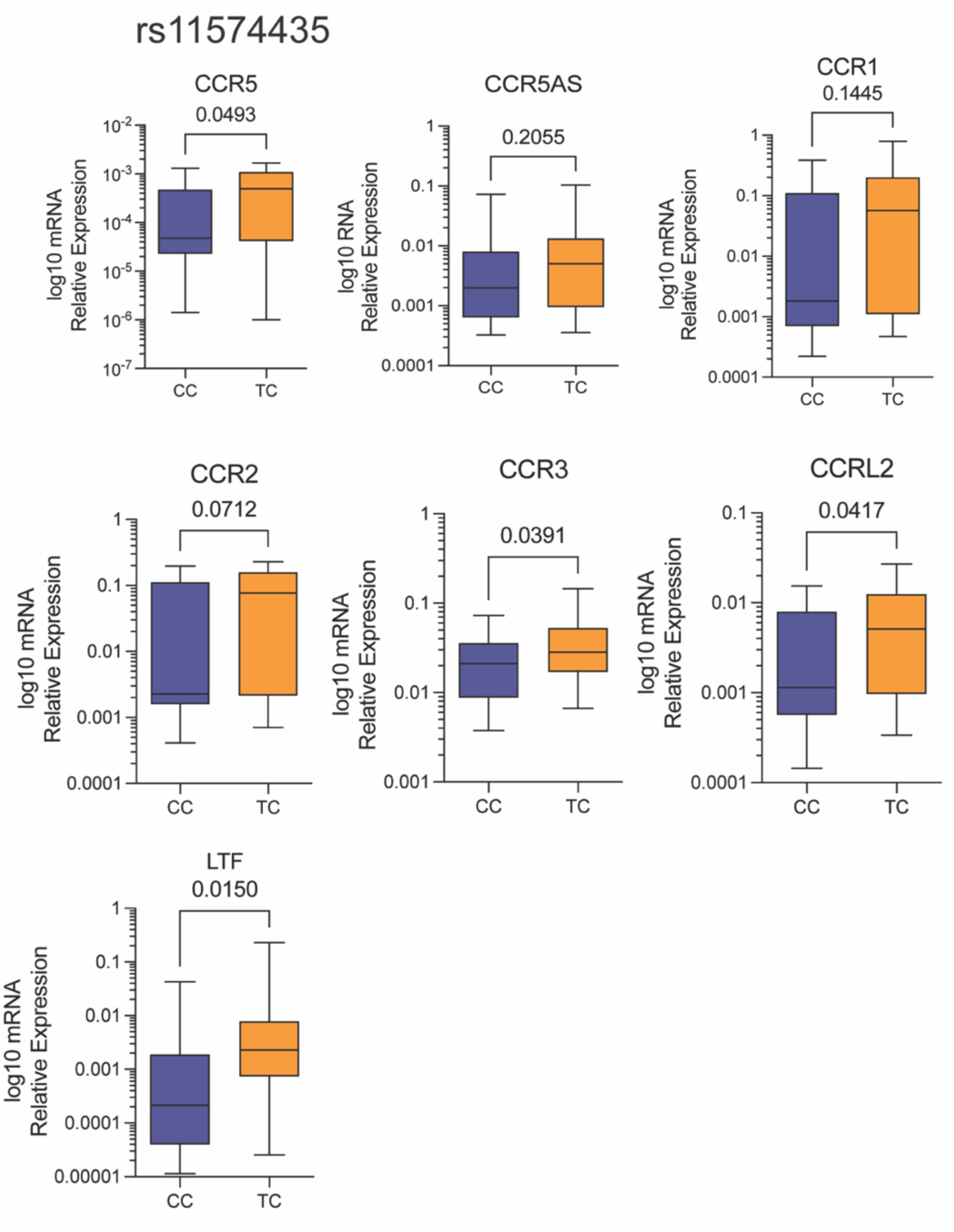
mRNA levels of *CCR1, CCR3, CCR2, CCRL2, LTF, CCR5* and *CCR5AS* were determined by RT-PCR and the values were stratified based on rs11574435. Data were analysed using Mann-Whitney U-test (P-value < 0.05).

**S11 Fig.**
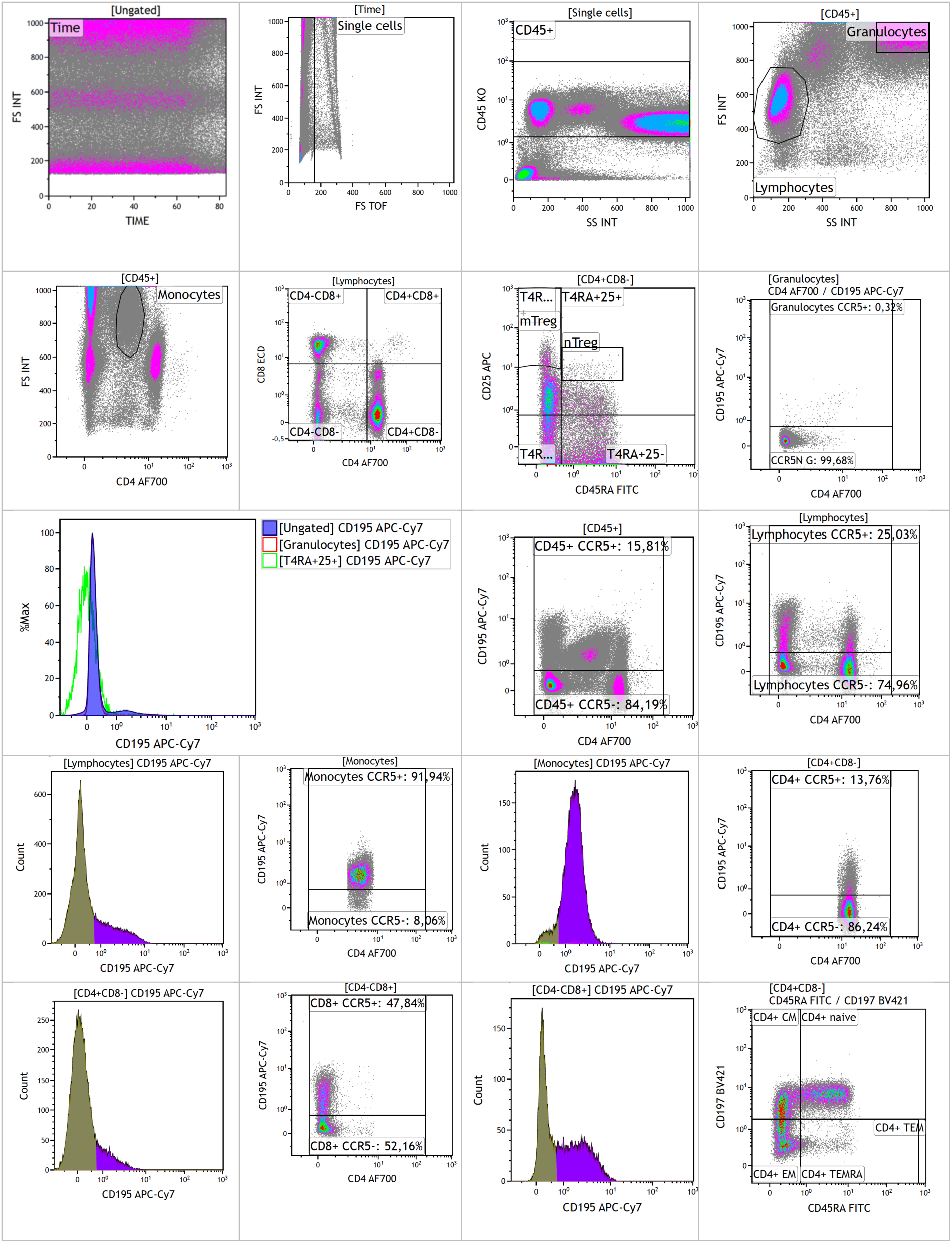
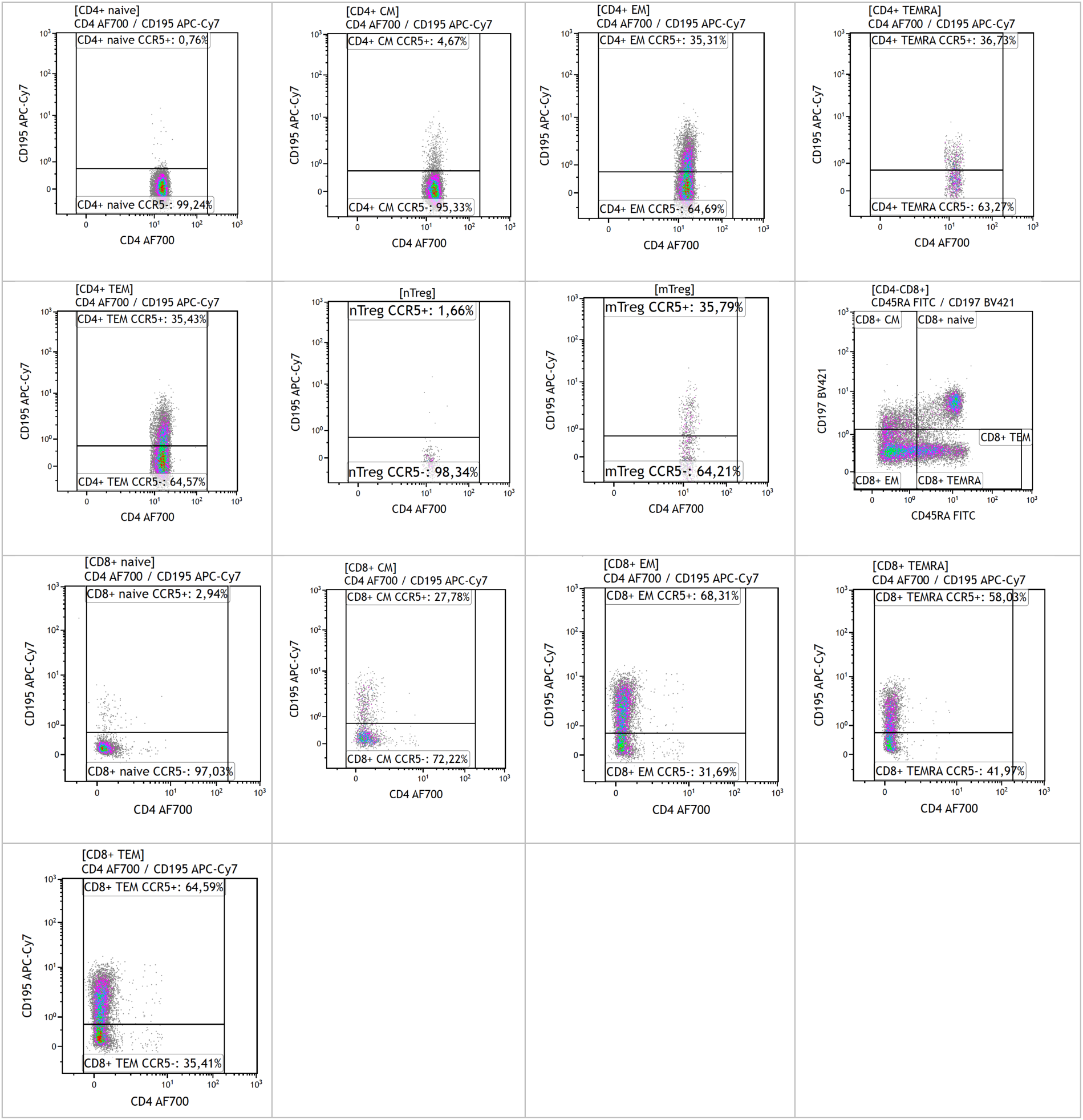
Example of the gating strategy. CD45+ cells were identified by gating on live and single cells and subsequently on CD45+ cells. Within the CD45+ cells, lymphocytes and monocytes were identified by granularity (side scatter) and size (forward scatter). Lymphocytes were further classified into different subsets of CD4+(CD8-) T cells and (CD4-) CD8+ T cells. CD4+ cells and CD8+ were classified as being naïve (CD45RA+CCR7+), central memory (CM, CD45RA-CCR7+), effector memory cells (EM, CD45RA-CCR7-), effector memory cells expressing CD45RA (TEMRA, CD45RA+CCR7-) and the total pool of effector memory cells (TEM, CD45RA-/+CCR7-). CD4+ naive regulatory (nTreg, CD45RA+CD25+) and CD4+ memory regulatory (mTreg, CD45RA-CD45++) cell subsets were identified within the subset of CD4+CD8-T cells.

### Methods

Live and single cells were selected first. Leukocytes were identified using CD45. Lymphocytes and monocytes were identified by granularity and size. Lymphocytes were further characterized using CD4, CD8, CD45RA and CCR7 to identify CD4+ cells and CD8+ naïve (CD45RA+CCR7+), central memory (CM, CD45RA-CCR7+), effector memory cells (EM, CD45RA-CCR7-), effector memory cells expressing CD45RA (TEMRA, CD45RA+CCR7-) and the total pool of effector memory cells (TEM, CD45RA-/+CCR7-) (41, 42). In addition, CD4+ naive regulatory (nTreg, CD45RA+CD25+) and CD4+ memory regulatory (mTreg, CD45RA-CD25++) cell subsets were identified. Gates for CCR5 were set using an internal negative control (granulocytes) and fluorescence minus one controls. The regions that identified CCR5-cell populations in granulocytes were used to distinguish between CCR5- and CCR5+ cell populations in other cell types as well. The percentage of CCR5+ cells (%) and CCR5 geometric mean fluorescence intensity (MFI) were assessed on all identified cell types.

**S3 Table.**
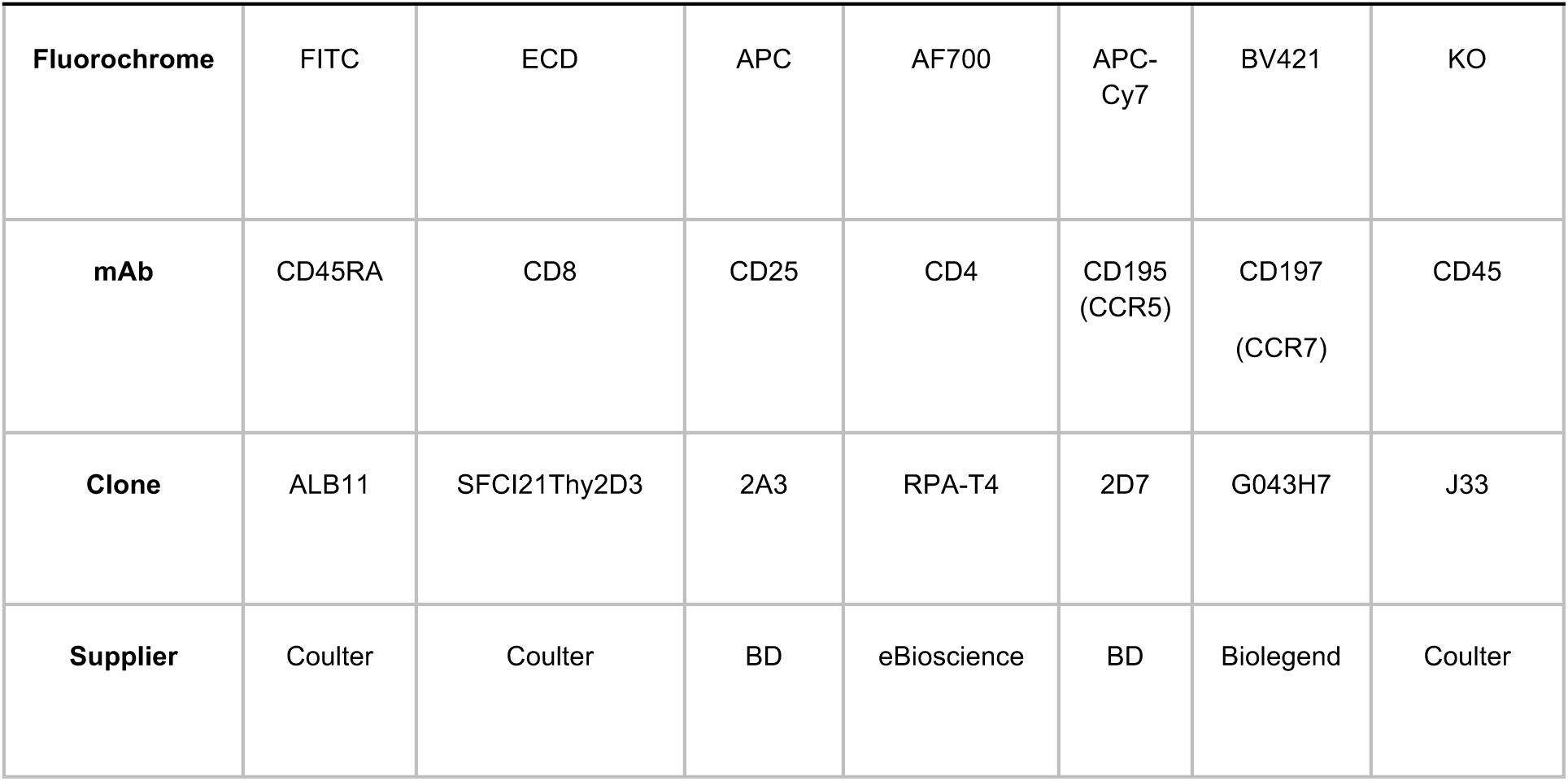
Summary of the antibody clones and the fluorochrome conjugates used for the fluorescent staining mixes. mAb = monoclonal antibody.

| Fluorochrome | FITC | ECD | APC | AF700 | APC-Cy7 | BV421 | KO |
| --- | --- | --- | --- | --- | --- | --- | --- |
| mAb | CD45RA | CD8 | CD25 | CD4 | CD195 (CCR5) | CD197 (CCR7) | CD45 |
| Clone | ALB11 | SFCI21Thy2D3 | 2A3 | RPA-T4 | 2D7 | G043H7 | J33 |
| Supplier | Coulter | Coulter | BD | eBioscience | BD | Biolegend | Coulter |

**S4 Table.**
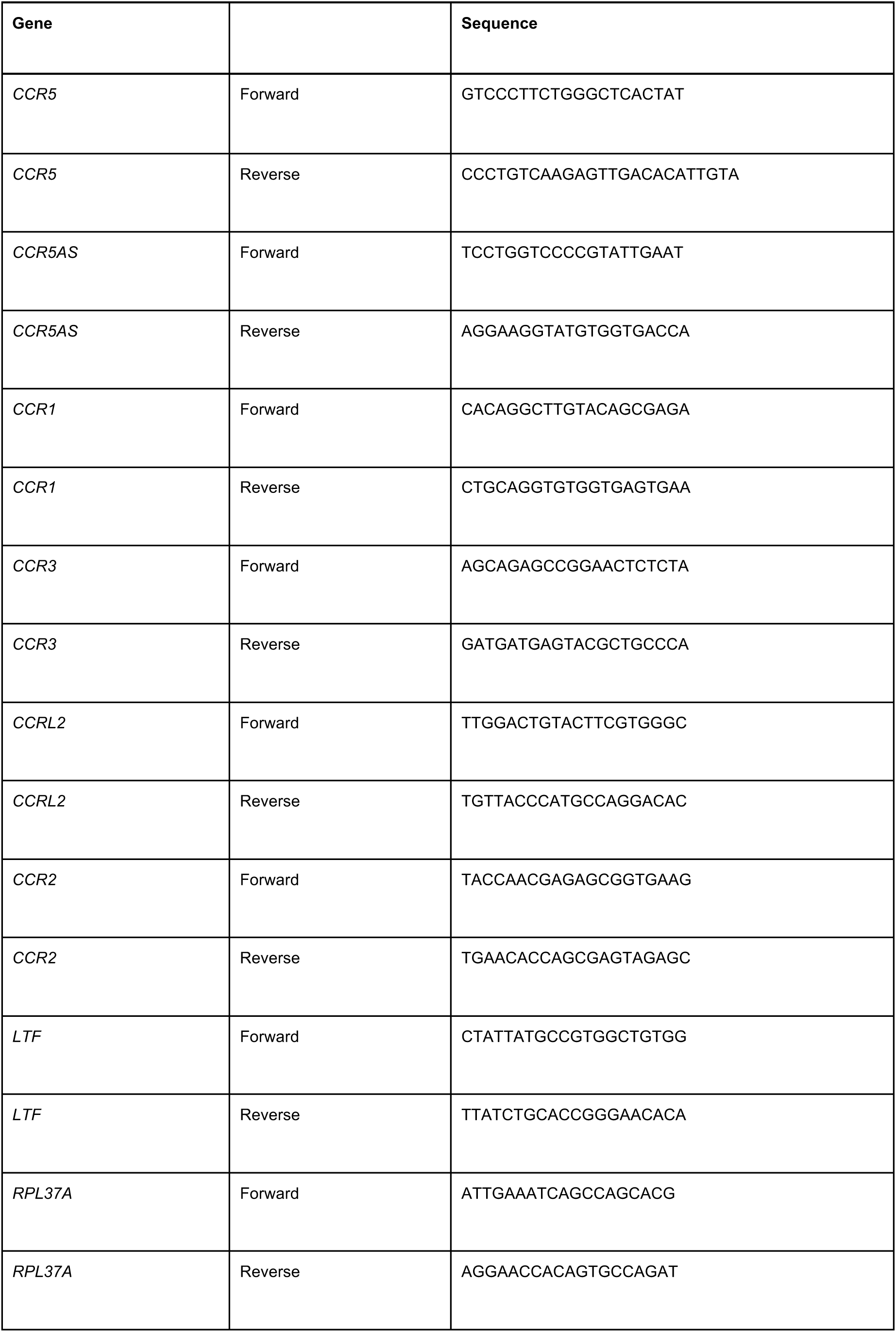
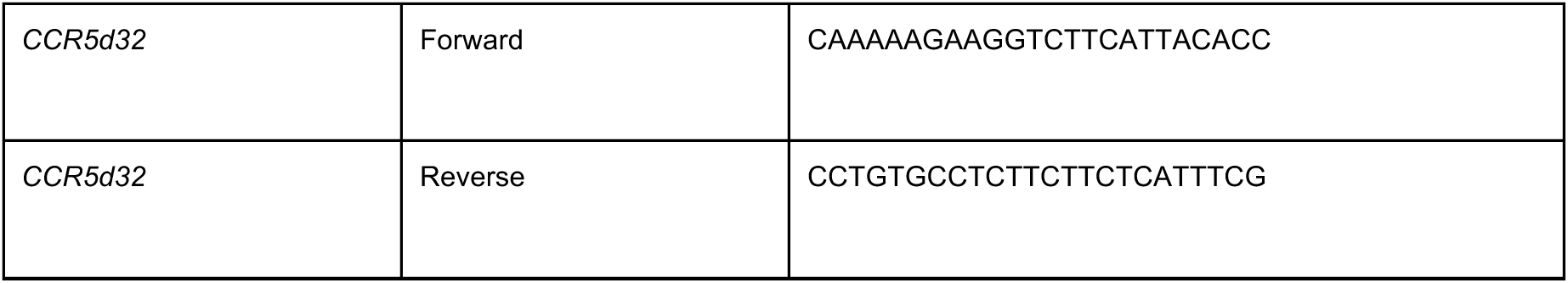
Primers sequences used in the qPCR and *CCR5d32* PCR

